# Constructing An Adult Orofacial Premotor Atlas In Allen Mouse CCF

**DOI:** 10.1101/2021.02.18.431923

**Authors:** Jun Takatoh, Jae Hong Park, Jinghao Lu, Shun Li, P. M. Thompson, Bao-Xia Han, Shengli Zhao, David Kleinfeld, Beth Friedman, Fan Wang

## Abstract

Premotor circuits in the brainstem control pools of orofacial motoneurons to execute essential functions such as drinking, eating, breathing, and in rodent, whisking. Previous transsynaptic tracing studies only mapped orofacial premotor circuits in neonatal mice but the adult circuits remain unknown due to technical difficulties. Here we developed a three-step monosynaptic transsynaptic tracing strategy to identify premotor neurons controlling whisker, tongue protrusion, and jaw-closing muscles in the adult. We registered these different groups of premotor neurons onto the Allen mouse brain common coordinate framework (CCF) and consequently generated a combined 3D orofacial premotor atlas, revealing unique spatial organizations of distinct premotor circuits. We also uncovered premotor neurons simultaneously innervating multiple motor nuclei and, thus, likely coordinating different muscles involved in the same orofacial behaviors. Our method for tracing adult premotor circuits and registering to Allen CCF is generally applicable and should facilitate the investigations of motor controls of diverse behaviors.

## Introduction

Orofacial behaviors, such as breathing, drinking, and eating, are essential for many animals to access vital sustenance (oxygen, water, and food). For example, water can be consumed by a consecutive orofacial sequence, including jaw opening, licking and swallowing. Mice, as nocturnal animals, use whisking and sniffing, often in coordination with breathing, to explore their physical environment (Deschenes et al., 2012; Kleinfeld and Deschenes, 2011; Moore et al., 2014). Thus, many of the orofacial behaviors utilize multiple muscles through coordinated activity of their associated motoneurons in a seamless manner (Kurnikova et al., 2017; McElvain et al., 2018; Moore et al., 2014). Given that each orofacial motoneuron pool only projects to its corresponding muscle and lacks collaterals to innervate other central neurons, coordinated orofacial behaviors are thought to be achieved in part by orofacial premotor circuits in the brainstem. Thus, to understand how these brainstem circuits integrate information of external sensory stimuli, self-motions, and animals’ needs or internal states to orchestrate the activities of multiple, distinct groups of cranial motoneurons, a key first step is to delineate the orofacial premotor circuit for each individual group of motoneurons in the adult nervous system. Putative premotor neurons projecting to different craniofacial motor nuclei had been previously mapped using conventional retrograde tracers injected into different motor nuclei in the brainstem (Aldes, 1990; Appenteng and Girdlestone, 1987; Borke et al., 1983; Chandler et al., 1990; Hattox et al., 2002; Isokawa-Akesson and Komisaruk, 1987; Li et al., 1996, 1997; Mizuno et al., 1983; Travers and Norgren, 1983; Vornov and Sutin, 1983). However, each orofacial motor nucleus contains functionally different and also antagonistic motor neurons (Aldes, 1995; Ashwell, 1982; Furutani et al., 2004; Gestreau et al., 2005; Hinrichsen and Watson, 1984; Klein and Rhoades, 1985; Komiyama et al., 1984; Krammer et al., 1979; Limwongse and DeSantis, 1977; Mizuno et al., 1975; Terashima et al., 1993; Watson et al., 1982). For instance, the hypoglossal nucleus contains motoneurons for tongue protrusion, retrusion, and shaping. Injection of conventional retrograde tracer into the hypoglossal nucleus would, therefore, label premotor circuits for all these groups of muscles (Aldes, 1995; Gestreau et al., 2005; Krammer et al., 1979). Moreover, injection of conventional tracer could label neuronal populations projecting to non-motoneurons (e.g. interneurons) adjacent to the injection site. Thus, a major limitation of conventional tracers is the lack of muscle/motoneuron specificity.

To overcome such limitations, we and others used a monosynaptic retrograde rabies virus transsynaptic tracing strategy and examined the premotor circuits for various craniofacial and somatic motoneurons in neonatal mice (Sreenivasan et al., 2015; Stanek et al., 2014; Stepien et al., 2010; Takatoh et al., 2013; Tripodi et al., 2011). In this strategy, the specific muscle of interest is inoculated with a glycoprotein (G protein)-deleted RV (ΔG-RV) encoding a fluorescent protein, and ΔG-RV is taken up by motor axon terminals in the muscle and retrogradely transported to the motoneuron cell bodies. ΔG-RV is trans-complemented with G protein in motor neurons and subsequently spread into premotor neurons. The lack of G protein in premotor neurons prevents further retrograde traveling of ΔG-RV (Figure 1A). However, ΔG-RV does not infect from peripheral muscles efficiently in animals older than P8, thus precluding the use of this strategy to trace premotor circuits beyond ∼ P15. Since many orofacial behaviors do not fully develop until after weaning (Westneat and Hall, 1992), it is uncertain whether the orofacial premotor circuits revealed for neonatal mice are the same in adult animals, when most of the behavior and electrophysiology experiments are conducted.

**Figure 1.**
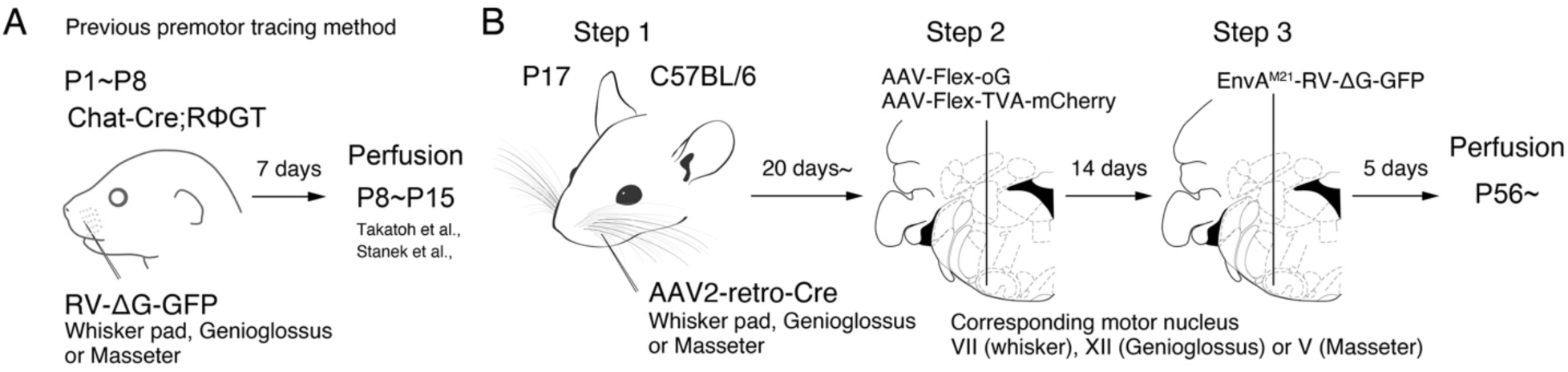
Monosynaptic rabies virus tracing strategy for labeling adult orofacial premotor circuits. (**A**) Schematic of previously used monosynaptic premotor transsynatptic tracing method in neonatal mice. (**B**) Schematic of the three-step monosynaptic premotor tracing strategy in adult mice developed in this study.

Another previously unsolved issue is to map traced premotor neurons from different muscles to a standard reference frame to allow cross-comparison of their spatial distributions. Earlier studies mapped and annotated the location of traced cells roughly based on outlines of brain sections compared to the standard stereotaxic atlas (Franklin and Paxinos, 2008). It is difficult to compare the spatial organizations for different tracing results, let alone comparing results across laboratories because many anatomical structures in brainstem are poorly defined. For example, the intermediate reticular nucleus (IRt), where many orofacial premotor neurons reside, extends ∼3 mm long in the adult along the anterior-posterior axis, thus simply annotating premotor neurons residing in IRt does not pinpoint the exact spatial location. Therefore, mapping premotor neurons in a standard coordinate frame and reconstructing them in the same 3-dimensional space with high spatial resolution would greatly facilitate subsequent functional interrogation of orofacial premotor circuits.

In the present study, we developed a new three-step monosynaptic rabies virus based strategy to trace orofacial premotor circuits in adult mice, and used this approach to delineate premotor circuits for whisker, genioglossus (tongue protrusion) and masseter (jaw-closing) motoneurons. The traced premotor neurons were all registered in the Allen Mouse Brain Common Coordinate Framework (CCFv3) (Wang et al., 2020). 3D reconstruction further enabled visualization of the full picture of their relative organization and distributions. The coordinates of all traced premotor neurons are accessible from the source file for interested users.

## Results

### Adult premotor circuit tracing strategy

To achieve monosynaptic orofacial premotor circuit tracing in adult mice, we developed a three-step monosynaptic rabies virus tracing strategy (Figure 1B). First, we introduce Cre recombinase into motoneurons innervating specific muscles through the intramuscular injection of the highly efficient retrograde viral vector AAV2retro-Cre (Tervo et al., 2016) in juvenile mice. Second, in adult mice, we inject Cre-dependent AAV expressing the TVA receptor and the optimized rabies glycoprotein oG (AAV-Flex-TVA-mCherry (Miyamichi et al., 2013) and AAV-Flex-oG (Kim et al., 2016)) into the corresponding brainstem motor nuclei in adult mice. In this way, TVA-mCherry and oG are specifically expressed in motoneurons that innervate the muscle that previously had AAV2retro-Cre injection. Finally, we inject the pseudotyped EnvA-ΔG-RV-GFP, which only infect TVA-expressing motoneurons. To further reduce any non-specific background infection by EnvA-ΔG-RV-GFP, we used a mutated version of the envelope, EnvA(M21), for pseudotyping -ΔG-RV. Thus EnvA(M21)-RV-ΔG-GFP was used for all our experiments (Sakurai et al., 2016). EnvA(M21)-ΔG-RV is also called CANE-ΔG-RV). Five days later, through the complementation of oG, the virus spread into the corresponding premotor neurons (Wickersham et al., 2007).

Initially, to determine the efficiency and specificity of the AAV2retro-Cre virus transduction of intended motoneurons, we injected the Cre virus into the whisker pad at different ages and examined the expression of Cre-dependent tdTomato (using Ai 14 mice) in the facial motor nucleus, where the myotopic map is well-described (Figure supplement 1) (Ashwell, 1982; Deschenes et al., 2016b; Furutani et al., 2004; Hinrichsen and Watson, 1984; Klein and Rhoades, 1985; Komiyama et al., 1984; Sreenivasan et al., 2015; Terashima et al., 1993; Watson et al., 1982). When we injected AAV2retro-Cre early in postnatal days (∼P10) into the whisker pad, we observed widespread labeling: in addition to motoneurons located in the lateral part of the facial nucleus (FN) that innervate the whisker pad, neurons in the medial and middle parts of FN were also labeled, likely due to more systemic infections. As the age advanced, the retrogradely labeled neurons became progressively restricted to the lateral part of FN, and the number of labeled neurons also decreased drastically (Figure supplement 1). We, therefore, decided to inject AAV2retro-Cre in the desired craniofacial muscles at P17 as step 1 which gave us specificity and good efficiency of infection. For step 2, at more than 3 weeks later, helper viruses (AAV-Flex-TVA-mCherry and AAV-Flex-oG) would be injected into the corresponding motor nucleus, and for step 3, 2 weeks after helper AAVs injected, EnvA(M21)-ΔG-RV-GFP would be injected into the same nucleus (Figure 1B). The brains are collected 5 days after RV injection.

We applied the three-step monosynaptic tracing strategy to investigate the premotor circuit for following motor units: whisker motoneurons (with cell bodies in lateral facial motor nucleus), tongue protruding genioglossus motoneurons of the hypoglossal nucleus, and the jaw-closing masseter motoneurons of the trigeminal motor nucleus, hereafter referred to as whisker, genioglossus, and masseter premotor circuits, respectively. The tracing results are described in details below. Notably, despite efficient transsynaptic labeling, we often did not observe TVA-mCherry and GFP double-positive motoneurons (starter cells). This loss of starter cells was due to the toxicity of ΔG-RV since omission of RV injection did not cause this problem (data not shown).

### Whisker premotor circuit

The densest labeling of whisker premotor neuron was found in: the Bötzinger complex (BötC)/retrofacial region (Figure 2A), the vibrissa zone of the IRt (vIRt) (Figure 2B), and the dorsal medullary reticular nucleus (MdD) (Figure 2E) (More quantitative analyses of the distribution of labeled cells in individual animals are described later in the paper). BötC/retrofacial region resides immediately posterior to the FN is known to contain expiration-rhythmic cells (Deschenes et al., 2016a), and is implicated in controlling sniffing behavior that is often coupled with whisking during exploration (Deschenes et al., 2012). vIRt, located medial to the compact part of the nucleus ambiguus (cNA), is known to contain whisking oscillator cells. A few premotor neurons were also consistently observed in the preBötinger complex (preBötC) (Figure 2B). We speculate that whisker premotor neurons in vIRt, BötC and preBötC are involved in modulating whisking rhythm.

**Figure 2.**
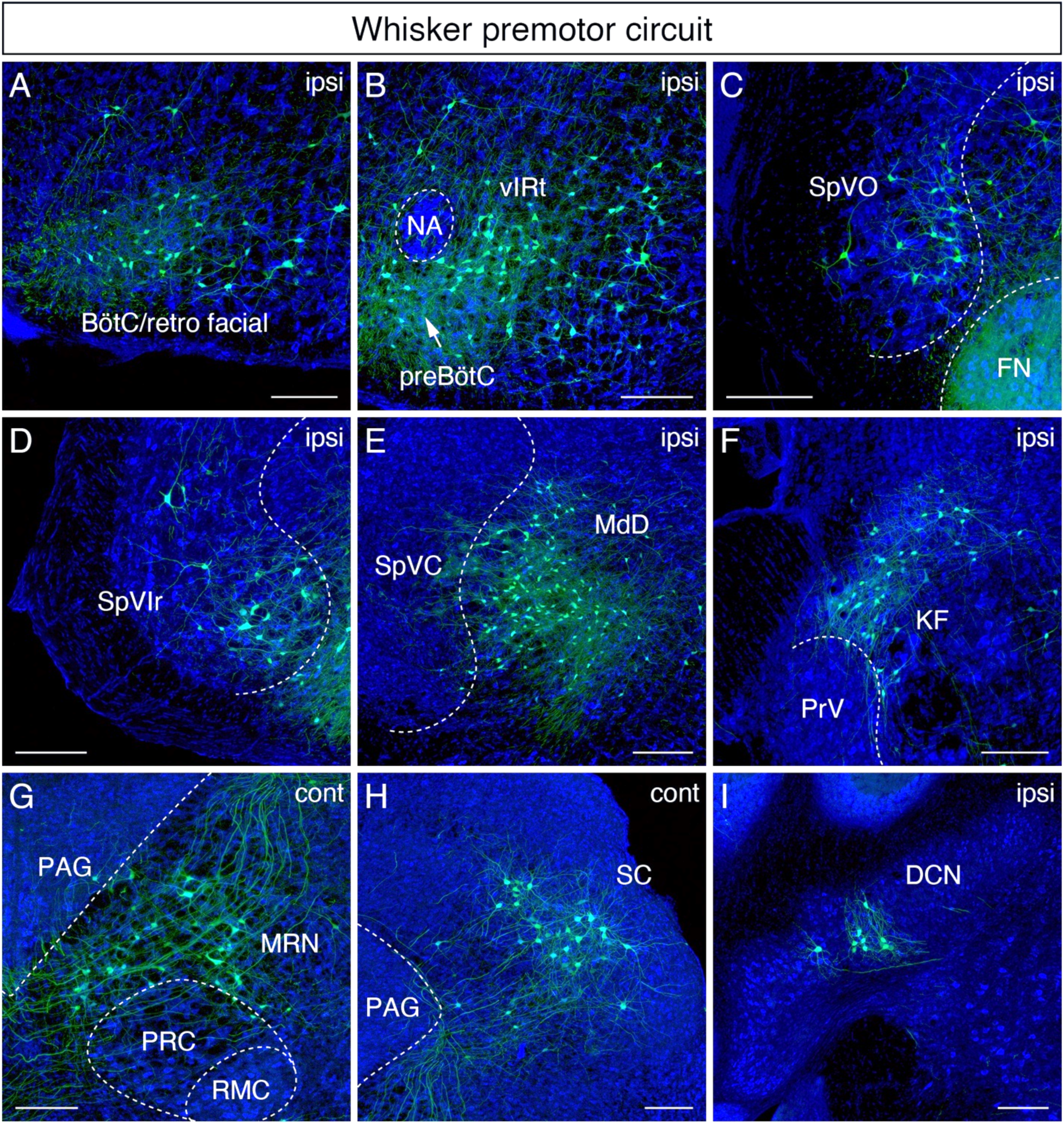
Monosynaptic tracing results of whisker premotor neurons in adult mice. Representative images of traced whisker premotor neurons on coronal sections. Sections were counterstained with fluorescent Nissl (blue). Labeled neurons shown in the ipsilateral BötC/retrofacial area (**A**), PreBötC (arrow) and vibrissal intermediate reticular formation (vIRt) (**B**), Spinal trigeminal nucleus oralis (SpVO) (**C**), rostral part of spinal trigeminal nucleus interpolaris (SpVIr) (**D**), medullary reticular nucleus dorsal (MdD) located medial to spinal trigeminal nucleus causalis (SpVC) (**E**), Kölliker-Fuse (KF) (**F**), contralateral midbrain reticular nucleus (MRN) located dorsal to the red nucleus parvicellular region (RPC) (**G**), contralateral superior colliculus (SC) (**H**), and ipsilateral deep cerebellar nucleus (DCN) (**I**). Scale bars, 200 µm.

We also found labeled neurons in the sensory-related areas, either within or adjacent to the ipsilateral spinal trigeminal nuclei, which receive inputs from whisker primary afferents. Those areas include the spinal trigeminal nucleus oralis (SpVO) (Figure 2C), rostral part of the interpolaris (SpVIr) (Figure 2D), and the muralis (SpVm, data not shown). Rostral to FN, we observed labeling in the ipsilateral Kölliker-Fuse (KF) (Figure 2F), the bilateral midbrain reticular formation (MRN, near the red nucleus) (Figure 2G), and the superior colliculus (SC) with contralateral dominance (Figure 2H). SC contains two clusters of whisker premotor neurons (Figure supplement 2A and 2B). The caudal cluster (Peak density; AP −3.73 ± 0.16 mm, ML 1.34 ± 0.03 mm, DV −2.57 ± 0.09 mm; n = 4 animals) resides in the intermediate layer of SC, whereas the rostral cluster locates in the deep layer of SC (Peak density; AP – 3.61 ± 0.09mm, ML 1.09 ± 0.04mm, DV −2.57 ± 0.09; n = 4 animals).

Note that while we aimed our injection at the intrinsic whisker muscles controlling whisker protraction, we could not rule out the possibility of infecting a few extrinsic motoneurons regulating whisker-pad retraction. We did not want to lesion the nerve innerving the extrinsic pad muscle in order to make the tracing more specific for intrinsic muscle, because the mice need to survive into adult (8-9 weeks old) in order to trace adult premotor circuit and lesioning in juvenile mice could cause compensatory changes in the circuits. The distribution of whisker premotor neurons observed in adult mice is consistent with the pattern observed previously in peri-natal tracing experiments (Takatoh et al., 2013). However, we did find a few differences between adult and postnatal circuits. First, whisker premotor neurons were labeled in the ipsilateral deep cerebellar nucleus interpositus (Figure 2I) in the adult mice, which was not present in juvenile animals. Second, we did not find premotor neurons in the spinal vestibular nucleus in adult mice, which was observed in peri-natal tracing. Third, the clusters of premotor cells in the lateral paragigantocellular nucleus (LPGi) bilaterally in neonatal mice became less distinct in adult. The neurons in LPGi might have migrated medially in post-juvenile development because we observed the larger number of labeled cells in the gigantocellular reticular nucleus, which situates medial to LPGi, at the level of the facial motor nucleus in adult mice. Finally, we observed labeled neurons in the zona incerta and in extended amygdala in adult that were not labeled in the neonatal transsynaptic tracing studies (Figure supplement 2C and C’) (Takatoh et al., 2013). Collectively, adult premotor tracing revealed both addition and loss of whisker premotor neurons in a few areas, and a similar spatial distribution patterns in the brainstem reticular and sensory nuclei in juvenile and adult mice.

### Tongue-protruding premotor circuit

We did not distinguish between the ipsilateral and contralateral results in the tongue premotor tracing since the left and right hypoglossal motor nucleus are adjacent to the midline and the genioglossus muscles are also located near midline such that the AAV2retro virus could infect motoneurons on both sides. The greatest number of labeled tongue-protruding premotor neurons was found bilaterally in the dorsal IRt (Figure 3A and 3B) (More quantitative analyses of the distribution of labeled premotor neurons in individual animals are described later in the paper). These neurons spread along the anterior-posterior axis of the dorsal IRt with the highest density in the area anterior to the rostral edge of the hypoglossal nucleus (see the details of IRt organization in the section below). Extending from the dorsal to ventral IRt (where vIRt resides), labeling gradually became sparser. Lateral and rostral to IRt, we also observed premotor neurons with relatively larger-sizes (compared to IRt) in the parvicellular reticular nucleus (PCRt) dorsal to the facial motor nucleus, and these cells exhibit medially oriented dendrites (Figure 3C). Many labeled cells with very large soma size were found in Gi/LPGi/LRt areas, spanning along the anterior-posterior axis (AP coordinate) (Figure 3D). In the pons, labeled premotor neurons were found in the supratrigeminal nucleus and peritrigeminal zone around the trigeminal motor nucleus (Figure 3E). In the sensory-related areas, labeled tongue premotor neurons were observed in the dorsal part of the principal trigeminal nucleus (PrV) (Figure 3C), nucleus solitary tract (NST), and dorsomedial SpV (DMSpV) (Figure 3F). In the cerebellum, labeled neurons resided in the medial subnucleus of DCN (Figure 3G). Additional premotor input was found in the raphe obscurus nucleus (Ro) (Figure 3H). The distribution of the adult genioglossus premotor neurons described above is similar to the pattern observed in juvenile mice (P8>P15 transsynaptic tracing)(Stanek et al., 2014). However, in adult mice, the large cluster of premotor neurons in the dorsal midbrain reticular formation (dMRf) previously found in juvenile animals was absent.

**Figure 3.**
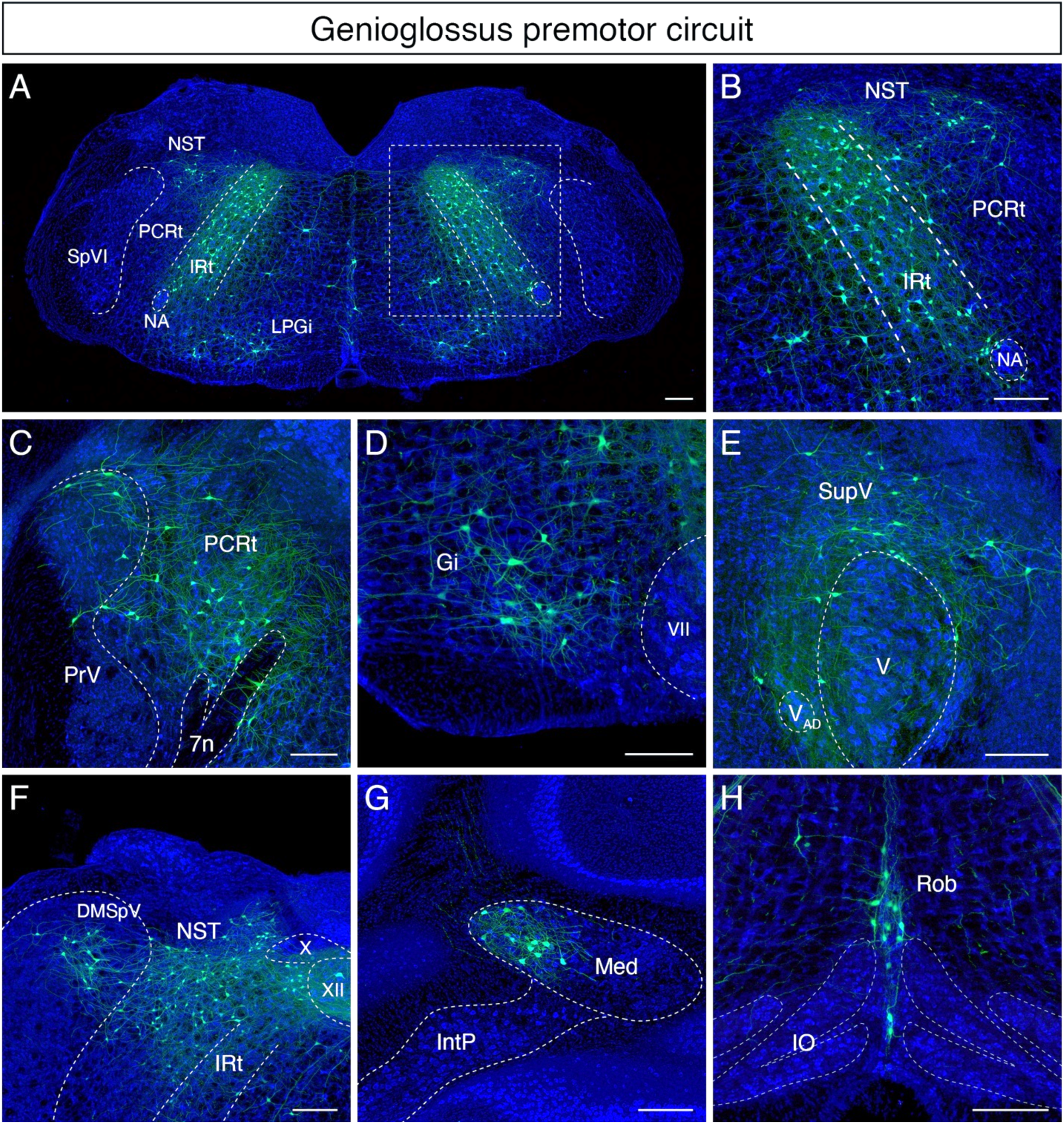
Monosynaptic tracing results of tongue-protruding genioglossus premotor neurons in adult mice. Representative images of traced genioglossus premotor neurons on coronal sections. Sections were counterstained with fluorescent Nissl (blue). Labeled neurons are shown in the dorsal intermediate reticular nucleus (IRt), nucleus of solitary tract (NST), lateral paragigantocellular nucleus (LPGi) at the anterior-posterior level between VII and XII (**A**, magnified view of the boxed area in **A** is shown in **B**), parvicellular reticular nucleus (PCRt), dorsal region of the principal trigeminal nucleus (PrV) (**C**), Gigangtocellular reticular nucleus (Gi) (**D**), supra-trigeminal region (SupV) (**E**), dorsomedial part of spinal trigeminal nucleus (DMSpV), rostral NST at the anterior-posterior level of the anterior part of XII (**F**), the medial subnucleus of the deep cerebellar nucleus (DCN) (**G**), and raphe obscurus nucleus (Rob) (**H**). Scale bars, 200 µm.

### Jaw-closing premotor circuit

In the masseter premotor circuit, extensive labeling was also found bilaterally along the anterior-posterior axis of the dorsal IRt (Figure 4A and 4B) (More quantitative analyses of the distribution of labeled masseter premotor cells in individual animals are described below). Interestingly, the majority of labeled dorsal IRt neurons were observed *contralaterally* in the caudal part of IRt (AP coordinate) (Figure 4C). Bilateral labeling in PCRt was observed as a lateral continuum of the dorsal IRt neurons at the level of the FN (Figure 4B). Rostrally, we found a distinct bilateral cluster of large-size neurons with medially directed dendrites situated around PCRt/PrV area immediately caudal to the trigeminal motor nucleus (Figure 4D). This group of neurons wedged into the dorsomedial and ventrolateral PrV. This area is identified by Nissl staining as containing a distinct cluster of neurons with large size than neighboring cells. Similar to the tongue-protruding circuit but with fewer numbers, cells of very large soma size were labeled ipsilaterally along the anterior-posterior axis spanning Gi/LPGi/LRt areas (AP coordinate) (Figure 4A). In the pons, numerous labeled masseter premotor neurons were also observed in the supratrigeminal nucleus and peritrigeminal areas (Figure 4D). In the sensory-related areas, labeled cells resided bilaterally in the dorsal PrV (Figure 4D), ipsilaterally in the dorsomedial SpV (Figure 4A), and ipsilaterally in the mesencephalic trig*eminal nucleus* (Figure 4E). In the cerebellum, we identified labeled neurons in the contralateral medial subnucleus of DCN (Figure 4F). The distribution of jaw-closing premotor neurons described above is similar to the pattern observed in juvenile mice (P8>P15 transsynaptic tracing) (Stanek et al., 2014). However, same as for the tongue premotor circuit, cells in dMRf observed in juvenile mice are absent in the adult circuit.

**Figure 4.**
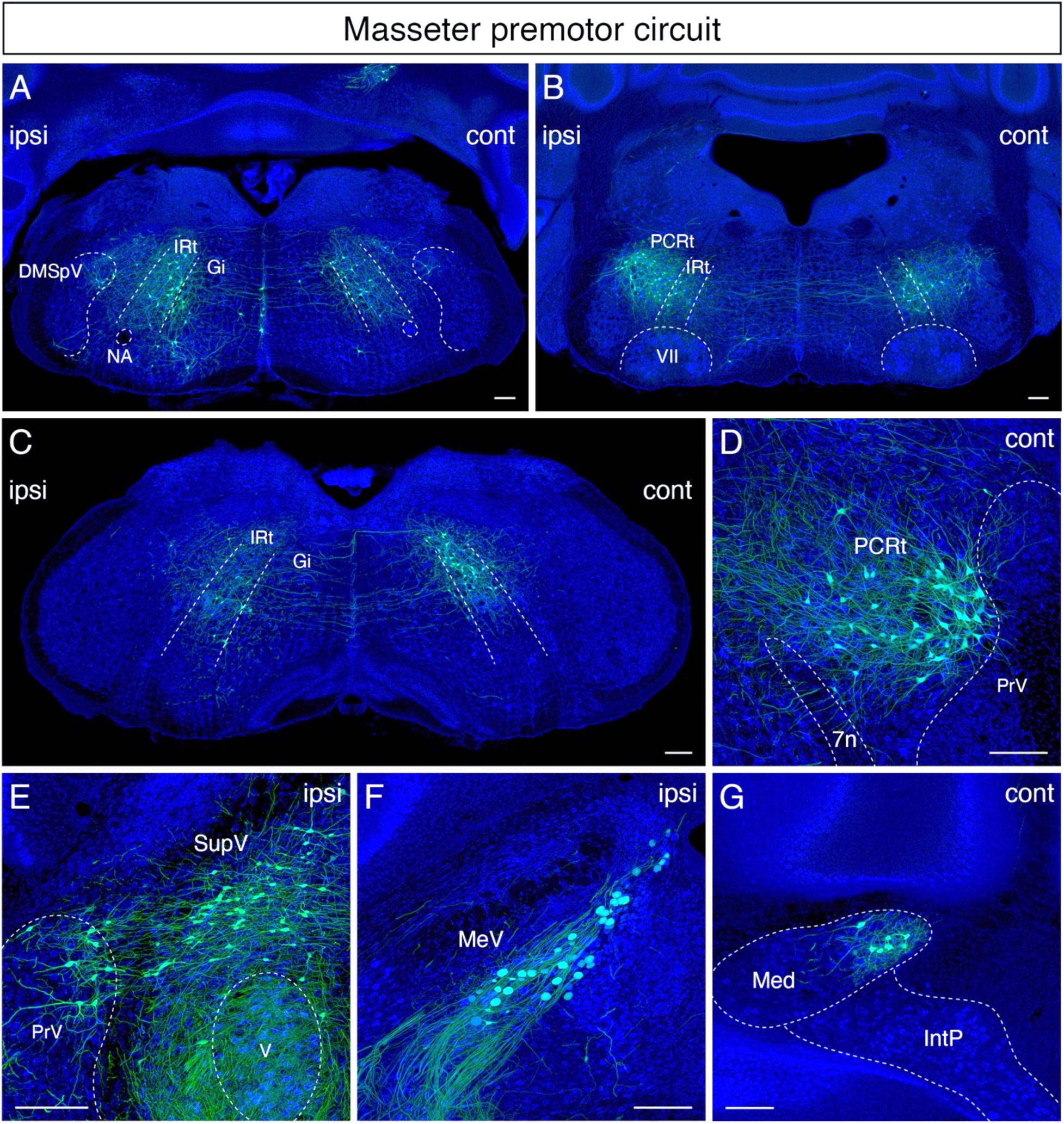
Monosynaptic tracing results of jaw-closing masseter premotor neurons in adult mice. Representative images of traced masseter premotor neurons on coronal sections. Sections were counterstained with fluorescent Nissl (blue). Labeled neurons are observed bilaterally in the the dorsal intermediate reticular nucleus (dorsal IRt), in dorsomedial part of spinal trigeminal nucleus DMSpV at the anterior-posterior level between VII and XII (**A**), bilaterally in the dorsal IRt, PCRt at the anterior-posterior level of VII (**B**), contralaterally in the dorsal IRt at the anterior-posterior level of the anterior part of XII (**C**), PCRt (**D**), SupV, dorsal PrV (**E**), ipsilateral mesencephalic nucleus MeV (**F**), PCRt, dorsal PrV (**C**), Gi (**D**), SupV (**E**) and the contralateral medial subnucleus of DCN (**G**). Scale bars, 200 µm.

### Mapping orofacial premotor neurons onto Allen common coordinate framework (CCF)

To generate standardized orofacial premotor atlas that enabling cross-comparison of different premotor circuits, RV-traced GFP-positive premotor neurons were mapped onto Allen Mouse Brain Common Coordinate Framework (CCFv3) (Wang et al., 2020). CCF is a widely used open-access 3D standardized brain atlas generated from the average of 1675 adult C57BL/6J mice. Registration of labeled neurons in CCF enables direct comparison of the results from different laboratories in the same coordinate space. Locations of RV-traced premotor neurons were translated into CCF coordinates using a method based on a previously described method with our modifications (Shamash et al., 2018). Briefly, each coronal section was registered to a corresponding CCF plane through diffeomorphic transformation (details see Methods). Subsequently, the labeled cells were identified and counted semi-automatically or manually, and their coordinates were transformed into CCF coordinates (Figure 5A). All traced orofacial motor neurons for whisker (n = 4 mice), genioglossus (n = 4 mice), and masseter (n = 4 mice) were registered to the CCF, and their coordinates are accessible from the source file. The cells in CCF coordinates were reconstructed in 2D and 3D spaces using Brainrender (Claudi et al., 2020).

**Figure 5.**
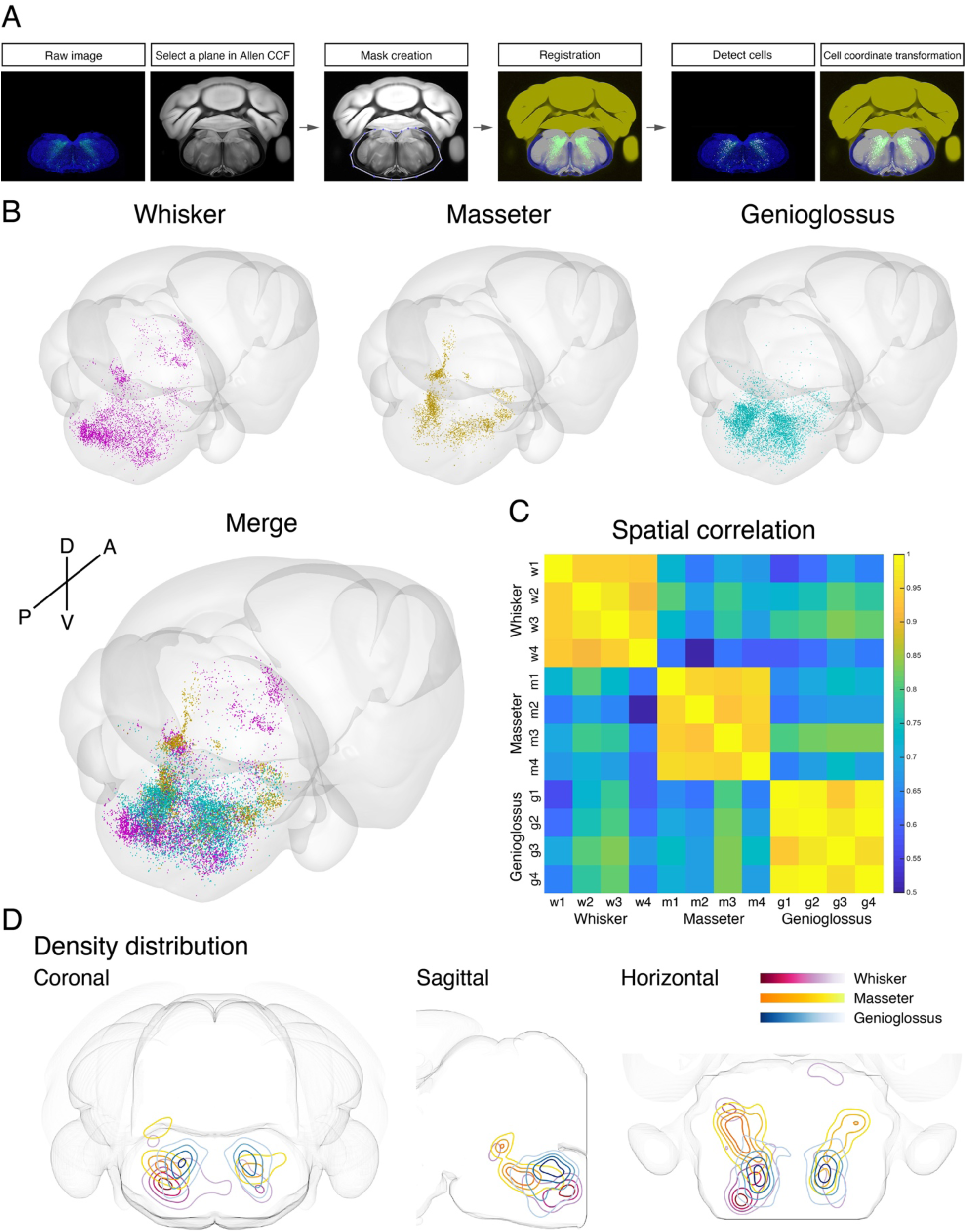
Co-registration and comparison of the spatial distributions of whisker, genioglossus, and masseter premotor circuits in Allen Common Coordinate Framework. (**A**) Procedure for mapping orofacial premotor neurons to Allen CCF. (**B**) Reconstructed representative whisker (magenta), masseter (gold), and genioglossus (cyan) premotor circuits in Allen CCF (top). Merged image (bottom). (**C**) Cross-correlation analysis of the spatial distribution patterns of individual animals. whisker (w1 - w4, n = 4), masseter (m1 - m4, n = 4), and genioglossus (g1 - g4, n = 4) premotor circuits. (**D**) 2D contour density analysis of representative whisker (magenta), masseter (yellow), and genioglossus (blue) premotor circuits.

### Cross-comparison of spatial distributions of whisker, tongue-protruding and jaw-closing premotor circuits

To compare the spatial organization of whisker, tongue-protruding, and jaw-closing premotor circuits, transsynaptic labeling results from individual animals were reconstructed in the same CCF space (Figure 5B, Supplemental Movie 1). Reconstructed premotor circuits for each of the target muscle/motoneurons from different animals are shown in Figure supplement 3-5. Using the extracted spatial coordinates of all labeled neurons, we performed cross-correlation analysis of the spatial distribution patterns of tracing results from all samples (see Methods). Individual premotor tracing results from the same muscle/motor unit were highly correlated, whereas results obtained from different muscles/motor units showed low correlations in spatial patterns (Figure 5C, Figure supplement 3-5). When we plot the premotor neurons for whisker/genioglossus/masseter into the same CCF in 3D, the results revealed both overlapping and segregating features of these different premotor circuits (Figure supplement 6, Supplemental Movie 1). The distribution density plot analysis of each premotor circuit also supports the muscle-specific differential spatial organizations as shown for all three planes: coronal, sagittal, and horizontal (Figure 5D). All three premotor circuits showed the highest density of labeling in the intermediate and parvicellular reticular formations (IRt and PCRt); however, the exact peak density positions were in the different locations of IRt/PCRt for different circuits. The whisker premotor circuit showed highest labeling density in the caudoventral areas of IRt (Figure 5D, Red). The masseter premotor circuit had densest labeling in the anterodorsal area of IRt (Figure 5D, Yellow). The highest-density area of the genioglossus premotor neurons located in IRt in between the peaks for the whisker and masseter premotor cells (along the A-P axis), although there were shared regions between genioglossus and masseter premotor distributions (Figure 5D, Blue). Finally, the extracted coordinates for each of the labeled cells enabled automatic assignment of their corresponding anatomical structure used by Allen CCF. Consequently, we can obtain the top 10 transsynaptically labeled premotor nuclei for each muscle recognized by the CCF (Figure Supplement 7). These analyses have collectively given us an overview of the differential anatomical organizations of whisker, tongue-protruding, and jaw-closing premotor circuits in adult mice.

### Detailed comparison of spatial Organization orofacial premotor circuits within IRt

Our adult tracing results indicates IRt as the common area containing premotor neurons for all three circuits. Since earlier studies have either localized or implicated IRt as the region containing oscillator neurons for several orofacial actions (i.e. whisking, licking, chewing rhythm), yet IRt is a poorly defined area, we decided to examine the relative spatial organizations of different premotor circuits within IRt in greater details. We took advantage of our reconstructions of whisker, genioglossus, and masseter IRt premotor neurons in the same CCF space to demarcate only cells within IRt (Figure 6, locations of craniofacial motor nuclei were also shown as landmarks). Again, these 3D reconstructions revealed partial overlapping and partial segregation of the three premotor circuits (Figure 6A–6C). Along the A-P axis, the highest density regions of ipsilateral jaw-closing and tongue-protruding premotor neurons in IRt were close to each other but with the peak of jaw premotor neurons shifted rostrally and ventrally (Figure 6G – 6O, jaw peak: AP −6.02 ± 0.18 mm, DV −6.45 ± 0.04 mm; n = 4 mice, tongue peak: AP −6.20 ± 0.23 mm, DV −5.74 ± 0.29; n = 4 mice). Along the D-V axis, while tongue premotor neurons are concentrated to more dorsal IRt than jaw premotor neurons, their distribution spread more to ventral IRt. Notably, the contralateral jaw IRt premotor neurons formed a discernable cluster caudal to the densest area of tongue IRt premotor neurons, displaying a bilaterally asymmetric distribution (Figure 6H). Whisker premotor neurons were more spatially separated from tongue and jaw premotor neurons in IRt, i.e at more caudal and ventral (AP −6.45 ± 0.19 mm, DV −6.45 ± 0.04 mm; n = 4 mice) locations in IRt (Figure 6D–6F). Furthermore, the tongue and jaw IRt premotor neurons showed similar densities between the ipsilateral and contralateral side (as licking and chewing generally involve muscles of both sides), the whisker IRt premotor neurons showed biased distribution to the ipsilateral side (Figure 6D and 6E). Collectively, these results suggest that functionally distinct groups of orofacial premotor neurons occupy the overlapping yet distinct spatial positions within IRt, and there is roughly a ventral-to-dorsal, and caudal-to-rostral gradient of whisker-tongue-jaw premotor neurons.

**Figure 6.**
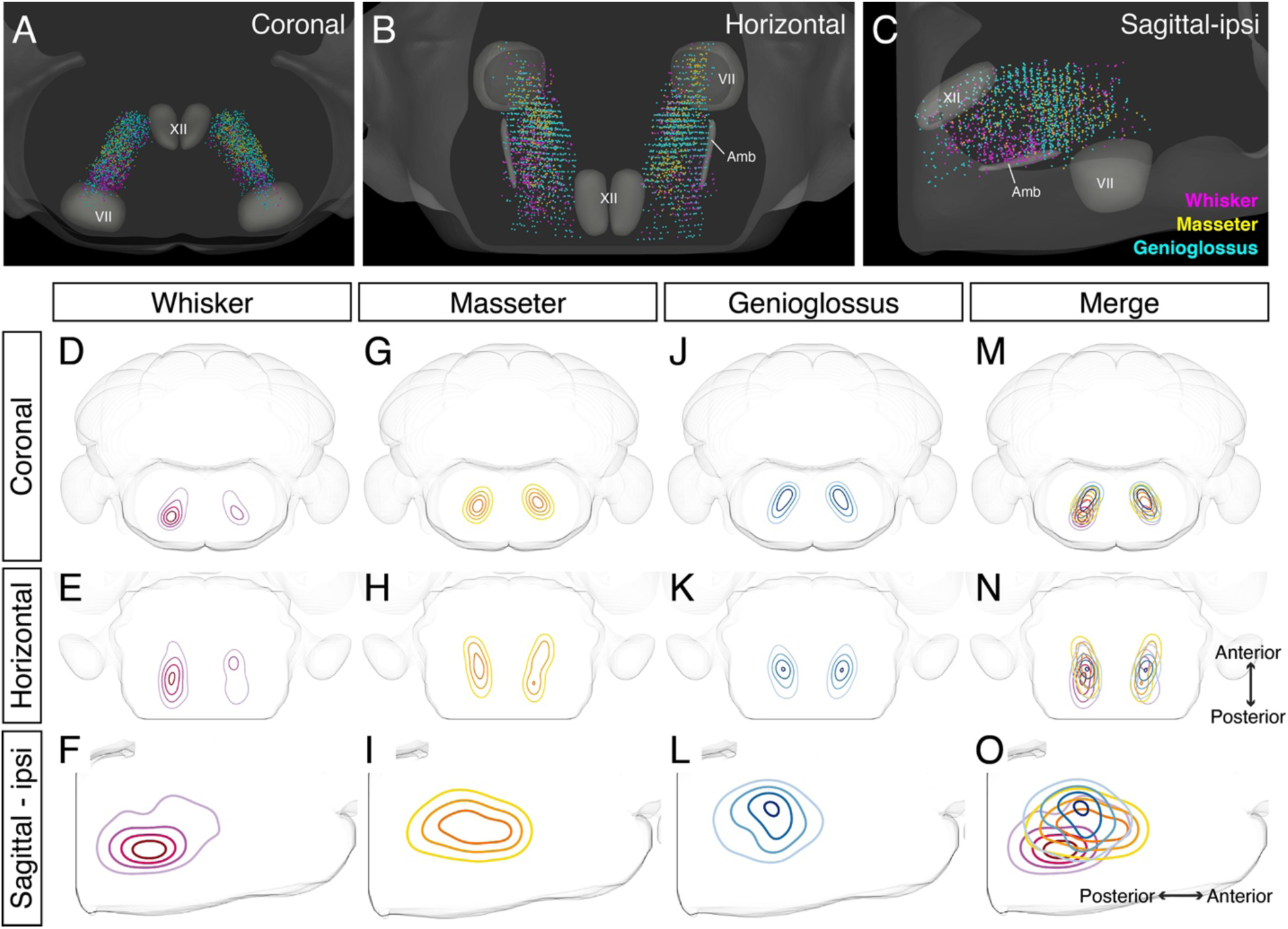
Detailed comparison of spatial organizations of orofacial premotor circuits within IRt. (**A-C**) Distribution of whiser (magenta), masseter (gold), and genioglossus (cyan) premotor neurons within IRt from representative animals in coronal (**A**), horizontal (**B**), and sagittal (**C**) planes. (**D-O**) Density analysis of whisker (**D-F**, magenta, an average of 4 mice), masseter (**G-I**, yellow, an average of 4 mice), genioglossus (**J-L**, blue, an average of 4 mice) premotor neuron distributions. Merged images (**M-O**).

### Axon collaterals revealed common premotor neurons for distinct motor neurons

As mentioned in introduction, orofacial behaviors often require coordinated activity of multiple groups of motoneurons. A premotor neuron that simultaneously innervates distinct motoneurons forms the simplest motor coordinating circuit. We therefore examined whether our premotor tracing results provide evidence for the existence of such common premotor neurons. Bright fluorescent signal from RV traced cells allows us to visualize their axons and collaterals. Interestingly, in genioglossus premotor tracing studies, axonal collaterals (from some labeled premotor neurons) were found in the middle part of the FN, VII_middle_ (Figure 7B), where motor neurons controlling lip and jaw (platysma) movements reside, and were also found densely innervating the small subnucleus of the trigeminal motor nucleus, V_AD_ (Figure 7C and 7C’), which controls the jaw-opening anterior digastric muscles, as well as were observed in the accessory facial motor nucleus (data not shown), which innervates the posterior digastric jaw-opening muscle. These results suggest that certain premotor neurons controlling tongue protrusion also simultaneously control mouth- and jaw-opening through their axon collaterals, providing a neural substrate for coordinating multiple motor groups needed for proper execution of behaviors such as licking and feeding.

**Figure 7.**
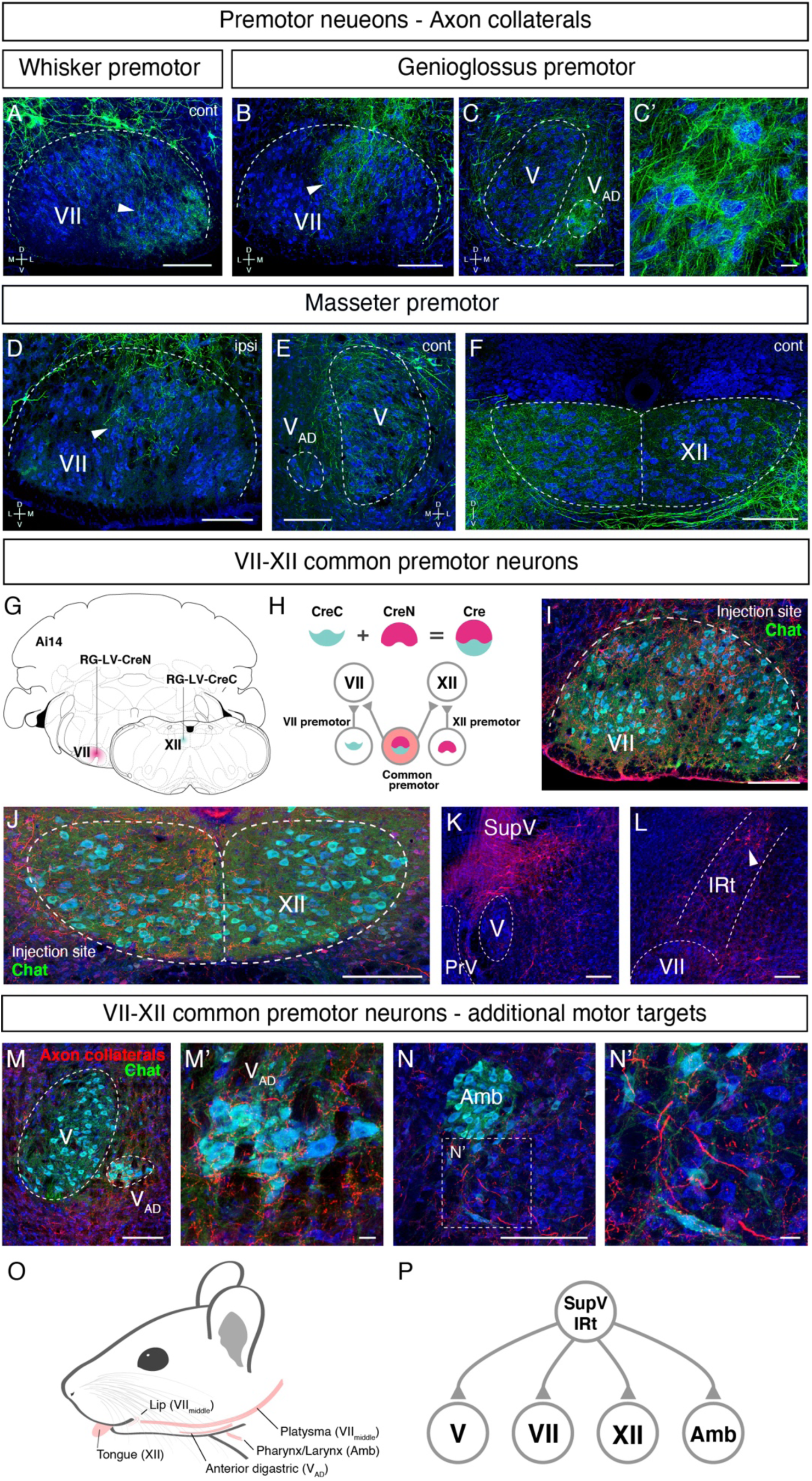
Common premotor neurons innervate multiple distinct orofacial motor nuclei. (**A-F**) Representative images of axon collaterals from rabies labeled premotor neurons traced from one muscle innervating other orofacial motor nuclei. Sections were counterstained with fluorescent Nissl (blue). (**A**) Axon collaterals from *ipsilateral* whisker premotor neurons innervate the *contralateral* whisker motoneurons in the lateral part of VII (arrowhead). (**B-C’**) Axon collaterals of some genioglossus premotor neurons also innervate the middle part of VII_middle_ (arrowhead, **B**), and innervate the anterior digastric part of V (V_AD_) (**C**, magnified view is shown in **C’**). (**D-F**) Axon collaterals from masseter premotor neurons also innervate the middle part of VII_middle_ (**D**), the contralateral V (**E**), and the dorsal part of XII (**F**). (**G-P**) Identifying VII_middle_-XII common premotor neurons. (**G, H**) Schematic of split-Cre tracing strategy. (**G**) RG-LV-CreN and RG-LV-CreC were injected into the left side of VII_middle_ and XII of Ai 14 mice, respectively. (**H**) Cre is reconstituted only in neurons innervating both VII_middle_ and XII, and which induces tdTomato reporter expression. (**I, J**) Representative images of axons/axon collaterals in the injection sites. Sections were counterstained with fluorescent Nissl (blue). Motoneurons were stained with anti-chat antibody (green). VII (**I**). XII (**J**). (**K, L**) Representative images of VII_middle_-XII common premotor neurons in SupV (**K**) and the dorsal IRt (**L**). (**M-N’**) Representative images of axon collaterals from VII_middle_-XII common premotor neurons in V_AD_ (**M**, magnified view in **M**’) and Amb (**N**, magnified view **N’**). Scale bars, 200 µm (**A-F**, **I-N**); 20 µm (**C’**, **M’**, **N’**). (**O**) Schematic showing orofacial muscle targets of motor nuclei. (**P**) Schematic of all motor nuclei innervated by VII_middle_-XII common premotor neuron in SupV and IRt.

Similarly, in masseter premotor tracing studies, the axonal collaterals of some labeled jaw-closing premotor neurons were observed in the middle part of the FN (Figure 7D) (VII_middle_, same region receiving innervations from the tongue premotor neurons), in the contralateral trigeminal motor nucleus (Figure 7E), and densely in the dorsal part of the hypoglossal nucleus (Figure 7F), where motor neurons for tongue-retrusion reside. In other words, the results suggest that premotor neurons controlling jaw-closing muscle also simultaneously modulate the tongue-retrusion and likely mouth closing muscles through their axon collaterals. In this manner, behaviorally synergistic motor units are coordinately activated to enable proper actions such as chewing without biting into the tongue. Finally, we also observed collaterals from whisker premotor neurons project to the contralateral later FN where whisker motoneurons reside (Figure 7A) (but not to hypoglossal and trigeminal motor nuclei), likely coordinating bilateral whisking.

Where might the common premotor neurons that send collaterals to multiple brainstem motor nuclei reside? Dense axon collaterals projecting to V_AD_ and VII_middle_ motoneurons from genioglossus premotor neurons inspired us to trace common premotor neurons innervating both XII (where motoneurons for genioglossus reside) and VII_middle_ using a retrograde split-Cre strategy. In this strategy, functionally inactive halves of Cre (CreN and CreC) packaged in retrograde lentivirus (RG-LV; RG-LV-CreN, RG-LV-CreC) were separately injected into VII_middle_ and XII of Ai14 mice (Figure 7G) (Stanek et al., 2016; Wang et al., 2012). In this injection scheme, functional Cre is reconstituted, and tdTomato is visualized only in neurons simultaneously innervating VII_middle_ and XII (Figure 7H). Retrograde split-Cre tracing revealed tdTomato-positive cells in SupV and the dorsal IRt areas (n = 4, Figure 7K and 7L), where genioglossus premotor neurons were found (Figure 3A, 3B, 3E and Figure supplement 4B). Notably, in addition to VII_middle_ and XII, we found tdTomato-positive axon terminals onto motoneurons in V_AD_ (jaw opening) and in the nucleus ambiguus (mostly in semi-compact part), which are known to be involved in swallowing. Thus, VII_middle_-XII common premotor neurons located in SupV and dorsal IRt simultaneously innervate motoneurons controlling tongue protrusion, lower lip, jaw-opening, and throat (through the nucleus ambiguus) (Figure 7O and 7P), suggesting that those common premotor neurons likely represent a fundamental neural substrate for coactivating these muscles. Interestingly, SupV and dorsal IRt were also labeled by the retrograde split-Cre tracing from the left and right sides of VII_middle_ (Figure supplement 8). Those neurons also project additional collaterals to V and XII, in addition to VII_middle_. These results indicate that SupV and dorsal IRt regions may be critical brainstem hubs containing common premotor neurons that coordinate multiple groups of motoneurons for orofacial feeding behaviors (Figure 7P).

## Discussion

We developed a three-step monosynaptic RV tracing to trace the premotor circuits for three different orofacial muscles in adult mice (Figure 8). We registered and reconstructed all the traced neurons in the standardized Allen mouse CCF and consequently generated the atlas showing positions of different orofacial premotor circuits in a common brain. This common atlas uncovers the overlapping yet distinct spatial organizations of premotor neurons involved in controlling movements of whiskers, the tongue and the jaw. Visualization of premotor neurons’ axon collaterals and retrograde split-Cre tracing studies further highlighted premotor neurons in SupV and dorsal IRt as potential substrates for coordinating multiple distinct orofacial muscles involved in feeding-related behaviors. Since these three groups of motoneurons are involved in three rhythmic orofacial behaviors, whisking, licking, or chewing, we next focus our discussion on the implications of the premotor atlas for rhythm generations.

**Figure 8.**
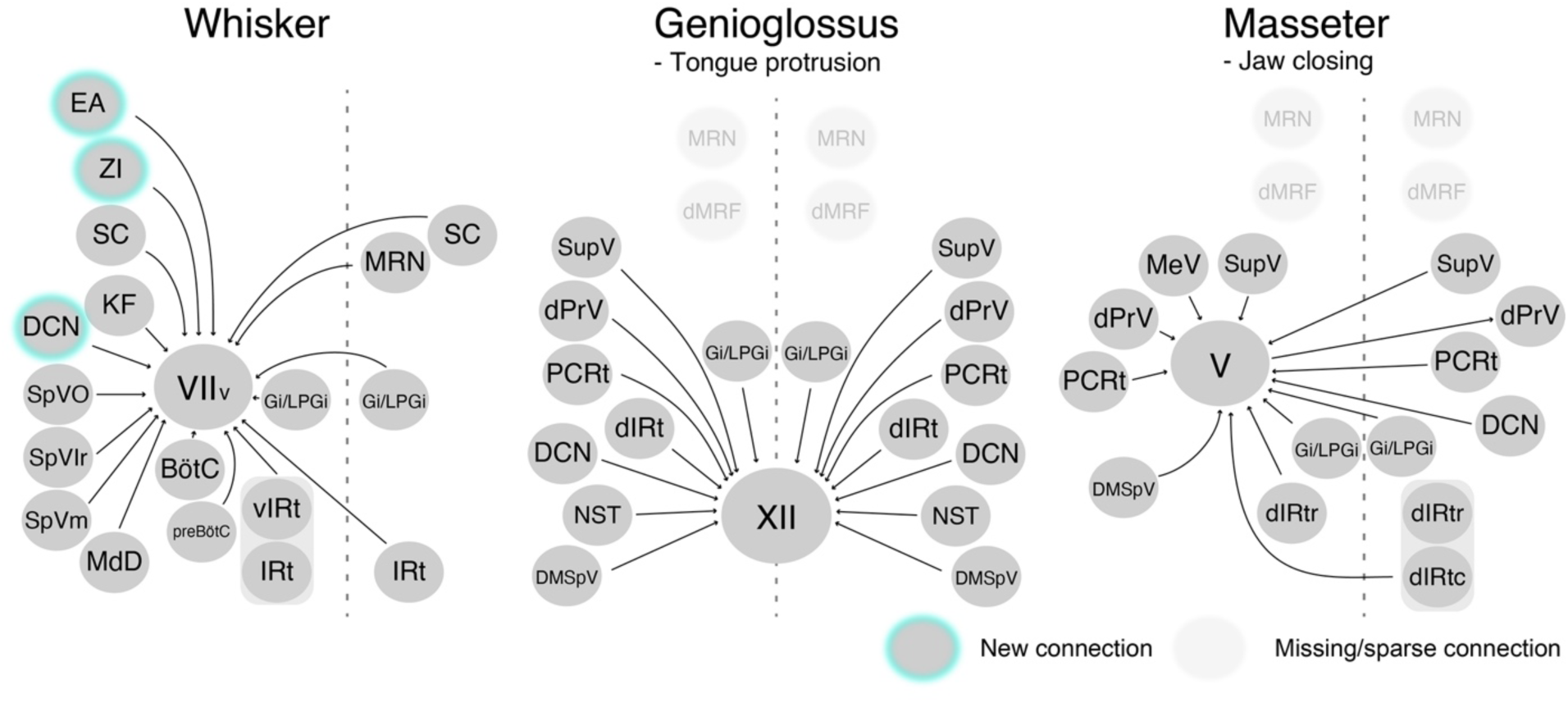
Schematic of whisker, tongue-protruding genioglossus, and jaw-closing premotor circuits in the adult mice. Newly emerged connections in adults that were not observed in neonates are outlined in turquoise. Neonatal connections that appear lost or becoming sparse are shown as translucent spheres.

### Implications for premotor neurons modulating whisking rhythm

Among the three orofacial premotor circuits in adult mice that we have mapped, the whisker premotor atlas consists of the most numerous brain structures (Figure 8). This is not surprising since dynamic whisker movements, as opposed to more stereotyped licking and chewing, are needed for tactile exploration of complex physical environment. Two previous studies uncovered vIRt as the region containing whisker oscillator neurons (Deschenes et al., 2016b; Moore et al., 2013). Indeed, we found extensive labeling of whisker premotor neurons in vIRt (with qualitatively more labeled neurons than neonatal tracing). Previous studies also revealed the coupling between breathing/sniffing and whisking (Moore et al., 2013; Welker, 1964). Along this line, we traced premotor cells in two brainstem areas known to control the respiratory rhythm, the retrofacial/BötC and preBötC (in both juvenile and adult mice), suggesting their roles in coordinating breathing and whisking, and potentially resetting whisking rhythm (Kleinfeld et al., 2014). However, there is an unresolved issue with regard to the inputs from the inspiratory rhythm generator preBötC. Moore et al showed that preBötC innervate vIRt, therefore preBötC is likely pre-premotor for whisker motoneurons (Moore et al., 2013). Deschênes et al. further demonstrated that in rats, a small injection of sindbis-GFP virus in electrophysiologically identified the lateral portion of preBötC revealed that these labeled preBötC neurons projects specifically to the lateral and dorsolateral part of FN where motor neurons for the nares dilation and extrinsic whisker-pad retraction reside, but rarely to the part where intrinsic whisker motoneurons locate (Deschenes et al., 2016b). In contrast, Yang and Feldman have shown that in mice, somatostatin- and glycine-positive prebötC neurons both project to the entire lateral part of FN, including ventral lateral FN where whisker-protracting intrinsic motoneurons reside (Yang and Feldman, 2018). In our tracing strategy, we injected AAV2retro-Cre into the whisker pad in P17 mice, therefore it is possible that we traced premotor neurons both for intrinsic and extrinsic motoneurons. Further study will be required to understand the precise connection between retrofacial/BötC and preBötC premotor neurons and extrinsic versus intrinsic motoneurons for whisking.

### Implications for premotor neurons modulating licking rhythm

In the tongue-protruding premotor circuit, dorsal IRt near the rostral end of the hypoglossal nucleus contains the highest density of premotor neurons. This area has previously been implicated as the rhythm generator for licking. Using extracellular recording in awake rats, Travers et al. demonstrated that neurons in this area show rhythmic activity phase-locked to licking (Travers et al., 2000). Furthermore, premotor neurons in this area express cFos after gaping behavior involving extensive tongue movement (DiNardo and Travers, 1997). However, bilateral infusion of muscimol in this area reduces licking EMG amplitude with minimal effect on the licking frequency (Chen et al., 2001; Travers et al., 2010), raising the possibility that dorsal IRt cells are the output of the actual licking oscillator. Importantly, bilateral infusion of muscimol in the same IRt area also suppresses chewing/mastication (see below), indicating the function of the dorsal IRt for coordination of tongue and jaw during ingestion behaviors. We also traced premotor neurons in NTS and the adjacent dorsolateral IRt. Interestingly, bilateral infusion of µ-opioid receptor agonist, Damgo, in the dorsolateral IRt/rostral NTS reduces licking frequency with increased amplitude. Future works using premotor neuron-specific manipulations will be necessary to dissect the function of the dorsal IRt and rostral NTS for modulating licking and chewing rhythms and amplitudes.

### Implications for premotor neurons modulating chewing rhythm

In the jaw-closing premotor circuit, two regions, the dorsal IRt and PCRt between the rostral extent of the FN and the rostral extent of the hypoglossal nucleus contain the highest numbers of premotor cells. This IRt/PCRt area has been implicated as a critical node for generating chewing rhythm (Chandler et al., 1990; Nakamura et al., 2017; Nozaki et al., 1986a, b; Travers et al., 2010). Nozaki et al., used a fictive rhythmic chewing preparation in guinea pigs to show that stimulation of the cortical masticatory area induces rhythmic activity in what was at the time called the oral part of the gigantocellular (Gi) reticular nucleus (Go). The rhythmic activity is, in turn, conveyed to the trigeminal motor nucleus through premotor neurons in the PCRt (Nozaki et al., 1986a, b). More recently, Travers et al., demonstrated in awake rats that muscimol infusion in rostral IRt/PCRt but not Go suppresses neuropeptide Y induced chewing behavior (Travers et al., 2010). Nakamura et al., used awake mice to show that infusion of bicuculline in the same IRt/PCRt area evokes chewing (Nakamura et al., 2017). Future studies with causal manipulation fo IRt/PCRt jaw premotor neurons should provide more definitive answers as to which neurons in IRt/PCRt and how are they involved in generating masticatory rhythms. Since there are also tongue-premotor neurons labeled in this area, it will be interesting to known whether these cells also innervate tongue-muscles and coordinating jaw-tongue movements during breaking down of food.

Other studies showed that neurons in the dorsal PrV generate rhythmic bursting in brainstem slices spontaneously and upon electrical stimulation of the trigeminal tract (Morquette et al., 2015; Sandler et al., 1998), and during fictive chewing in anaesthetized and paralyzed rabbits (Tsuboi et al., 2003), suggesting the role of this area for chewing rhythm generation. We also observed jaw premotor neurons in the lateral edge of PCRt (which could be part of SpVO or PrV) with an A-P location at the caudal edge of the trigeminal motor nucleus. Several studies assigned this area as a part of SpVO based on its receptive field in the oral areas (Inoue et al., 1992; Westberg et al., 1995; Yoshida et al., 1994). Neurons in this area in cats respond to either noxious stimulation of the tongue or to light mechanical stimulation of intra- or perioral structures, including the teeth, gingiva, and lip. These neurons issue collaterals that terminate in the trigeminal motor nucleus (Yoshida et al., 1994). Inoue et al., demonstrated in rats that premotor neurons in SpVO and the supratrigeminal nucleus share the same masticatory rhythm during cortically induced fictive mastication (Inoue et al., 1992). Some of these neurons are activated at short latencies by the stimulation of cortical masticatory area, or stimulation of the inferior alveolar and infraorbital nerves innervating oral areas, or passive jaw-opening. Based on these properties, it is suggested that premotor neurons in SpVO integrate sensory information from the oral area and rhythmic activity generated by a central rhythm generator to produce appropriate activity patterns during mastication.

### Common premotor neurons as potential neuronal substrate for coordinating orofacial behaviors

Tracing of axon collaterals from ΔG-RV-GFP labeled orofacial premotor neurons and the retrograde split-Cre mediated tracing studies uncovered common premotor neurons with extensive axon collateral network to jaw (trigeminal motor), tongue (hypoglossus), lip-jaw (facial) and throat (nucleus ambiguus), but importantly not to whiskers. The retrograde split-Cre tracing revealed that the major sources of these common premotor neurons are SupV and the dorsal IRt. Notably, these areas also contain bilaterally projecting masseter premotor neurons (Stanek). Neurons in both dorsal IRt and SupV are known to show rhythmic firing during licking (see Discussion above) and chewing (Inoue 1992). Common premotor neurons in SupV and dorsal IRt, therefore, may serve the simplest form of neuronal substrate for coordinating feeding-related orofacial behaviors by mean of broadcasting rhythmic information of licking and chewing to synergistic muscles. Future functional manipulation studies are needed to determine their causal functions in coordination of different orofacial behaviors.

### Other notable implications of the adult orofacial atlas

The three-step monosynaptic RV tracing allows us to reveal adult premotor circuits for a specific group of motor neurons, thereby advancing the transsyanptic premotor maps previously only available for neonatal mice. The coordinates of all traced neurons registered to the Allen mouse CCF are accessible from the source file, and can be used in the future to guide placement of electrodes for in vivo recordings as mice perform different orofacial behaviors. Comparing to neonatal circuits, we observed new additions of presynaptic inputs to control whisker motoneurons from ZI, DCN, and extended amygdala, and loss of presynaptic inputs from dMRF to tongue and jaw motoneurons in the adult premotor circuits (Figure 8). These changes may reflect more fine-tuned control of tactile whiskers, and changes in feeding behaviors from neonatal suckling to adult licking and chewing. Future studies with multi-color RV tracing using additional sets of recombinase and receptor-virus envelope, such as AAV2retro-FlpO, TVB-EnvB (Matsuyama et al., 2015) will enable simultaneous tracing of premotor circuits from two different motoneuron groups involved in the same orofacial actions. We envision transsynaptic premotor circuit tracing combined with functional characterization and activity manipulations will greatly advance our understanding of orofacial motor control.

## Materials and methods

### Animals

All animal experiments were conducted according to protocols approved by The Duke University Institutional Animal Care and Use Committee. Male and female C57B/L6 and Gt(Rosa)26Sor^tm14(CAG-tdTomato)Hze/J^ (Ai14) ( JAX # 007914) mice were obtained from the Jackson Laboratory (Bar Harbor, ME, USA) and used for virus tracing experiments.

### Viruses

AAV2retro-CAG-Cre

AAV2/8-CAG-Flex-TVA-mCherry (#48332, addgene Cambridge, MA, USA) (Miyamichi et al., 2013)

AAV2/8-CAG-Flex-oG (#74292, addgene) (Kim et al., 2016)

EnvA(M21)-RV-ΔG-GFP (also called CANE-ΔG-RV) (Sakurai et al., 2016; Wickersham et al., 2007)

### Monosynaptic transsynaptic rabies virus tracing

The tracing was performed in three steps.

### Peripheral tissue injection

To label a specific group of orofacial motor neurons, AAV2-retro-CAG-Cre (1000 µl, Harvard University, Boston Children’s Hospital Viral Core) was injected into either the whisker pad, genioglossus or masseter muscles at postnatal day 17, using a volumetric injection system (based on a single-axis oil hydraulic micromanipulator MO-10, Narishige International USA, Inc., East Meadow, NY, USA) (Petreanu et al., 2009) equipped with a pulled and beveled glass pipette (Drummond, 5-000-2005). Before injection, mice were anesthetized by a cocktail fo ketamine and xylazine (100 mg/kg and 10 mg/kg, i.p.). For the whisker pad, the virus was injected subcutaneously into the areas around C2 and B2 whiskers (500 nl each). For the genioglossus, the virus was injected directory into the muscle after exposing it by ventral neck dissection. Briefly, the genioglossus muscle was exposed by making a small incision in the mylohyoid muscle after the anterior digastric muscle was split open in the midline. For the masseter, the virus was injected into the area between the buccal and marginal nerves after making a small incision on a skin.

### Helper virus injection

For specific infection and glycoprotein complementation of pseudotyped RV-ΔG, helper viruses (120 nl, 1:1 mixture of AAV2/8-CAG-Flex-TVA-mCherry and AAV2/8-CAG-Flex-oG) were stereotaxically injected into the lateral part of the facial motor, hypoglossus, or trigeminal motor nuclei using a stereotaxic instrument (Model 963, David Kopf Instruments, Tujunga, CA, USA) three weeks or longer after the peripheral tissue injection. The viruses were injected at the rate of 30 nl/min with the injection system described above. The stereotaxic coordinates used were for the lateral part of the facial motor nucleus: 5.8 mm posterior, 1.38 mm lateral to the bregma, and 5.2 mm below the brain surface; for the hypoglossus nucleus: 5.8 mm posterior, 0.05 mm lateral to the bregma, and 5.15 mm below the brain surface with an anteroposterior 20° angle from vertical; and for the trigeminal motor nucleus: **4.1** mm posterior, 1.27 mm lateral to the bregma, and 4.6 mm below the brain surface with an anteroposterior 20° angle from vertical. Before suturing the skin, the craniotomy was filled with kwik-sil (World Precision Instruments, Inc., Sarasota, FL, USA) and covered with cyanoacrylate glue (Super Glue, Loctite, Westlake, Ohio, USA).

### Pseudotyped RV injection

Two weeks after the helper virus injection, EnvA(M21)-RV-ΔG-GFP (250 nl) was stereotaxically injected into the lateral part of the facial motor, hypoglossus, or trigeminal motor nuclei as described above.

### Retrograde split-Cre tracing

To label premotor neurons innervating multiple distinct motor nuclei, retrograde lentivirus carrying CreC or CreN (RG-LV-CreN and RG-LV-CreC) was stereotaxically injected separately into target motor nuclei of Cre-dependent tdTomato reporter mice. Specifically, for VII_middle_-XII premotor neurons, RG-LV-CreN (750nl) and RG-LV-CreC (500nl) were injected into VII_middle_ (5.8 mm posterior, 1.3 mm lateral to the bregma, and 5.2 mm below the brain surface) and the hypoglossus nucleus, respectively. For, bilateral VII_middle_ premotor neurons, RG-LV-CreN (750nl) and RG-LV-CreC (500nl) were injected into left and right VII_middle_, respectively.

### Histology

Five days after the pseudotyped RV injection, the animals were deeply anesthetized with isoflurane and transcardially perfused with 10% sucrose in Milli-Q water, followed by ice-cold 4% paraformaldehyde in 0.1 M phosphate buffer, pH 7.4. After dissection, the brains were post-fixed in the same fixative for overnight at 4°C and freeze-protected in 30% sucrose in phosphate buffer saline (PBS) at 4°C until they sank. The brains were embedded in OCT compound (Sakura Finetek USA, Inc., Torrance, CA, USA) and frozen in dry-ice-cooled ethanol. Eighty µm free-floating coronal sections were made using a cryostat (Leica Biosystems Inc, Buffalo Grove, IL, USA). The sections were briefly washed in PBS and stained with Neurotrace blue fluorescent Nissl stain (1:500, Thermo Fisher Scientific, Waltham, MA, USA) in 0.3% Triton-X100/PBS for overnight at 4°C. The sections were briefly washed and mounted on slide glasses with Mowiol. For retrograde split-Cre tracing experiments, some sections were stained with rabbit anti-RFP (1:500, #600-401-379, Rockland Immunochemicals, Inc. Limerick, PA, USA) and goat anti-choline Acetyltransferase (1:500, #AB144P, MilliporeSigma, Burlington, MA, USA) antibodies. Primary antibodies were visualized using donkey anti-rabbit antibody conjugated with Alexa Fluor Plus 555 (1:1000, #A32794, Thermo Fisher Scientific) and donkey anti-goat antibody conjugated with Alexa Fluor Plus 488 (1:1000, #A32814, Thermo Fisher Scientific).

### Imaging

Fluorescent images for atlas registration were taken with a Zeiss 700 laser scanning confocal microscope (Carl Zeiss Inc., Thornwood, NY, USA) using a 10x objective (pixel size, 1.042 × 1.042 µm).

### Mapping of Labeled neurons in Allen common coordinate framework

A previously published method SHARP-track (Shamash et al., 2018) was modified to improve registration of the brainstem sections to Allen CCF. Briefly, three steps for the registration were either introduced or improved, including (i) user-assist nonrigid deformation registration, (ii) correct conversion of coordinates (ML and DV to Bregma) after diffeomorphic registering of a section to the Allen CCF along the AP axis, and (iii) an option for automatic cell identification. First, the affine transform used by SHARP-track for brain section to reference registration is upgraded to LogDemons methods (Lu et al., 2018), a fast diffeomorphic registration method that can handle a more diverse scenarios of section distortion. Second, the procedures to determine the coordinates of the brain sections after nonrigid transformation and registration were corrected, and this is also critical for 3D reconstruction and visualizations of the results from serial 2D sections. Third, in addition to the manual cell identification by user generated click, an optional automatic cell identification function was developed to recover most identifiable cells, and subsequently users can manually correct mistakes. The automatic cell identification method contains a series of simple filters which balanced the speed and the precision of the approximation. Detailed implementation can be found in the Github repository (https://github.com/wanglab-neuro/Allen_CCF_reg). This site will be freely available upon publication. Coordinates of Bregma in Allen CCF was set at AP, 5400; ML, 5700; DV 0 (Shamash et al., 2018).

### Spatial correlation analysis

All 3D coordinates of the identified cells per mouse were concatenated. The spatial distribution was then estimated using multivariate kernel smoothing density function estimation. The estimated multivariate (3D) density functions for each mouse were then vectorized and pairwise cosine similarity was computed for all the mice. The result was shaped to a square matrix which was shown in the figure.

### Visualization of labeled neurons on Allen common coordinate framework

Premotor neurons registered in Allen CCF were visualized using Brainrender (Claudi et al., 2020) or a custom written code (for density plots). Briefly, the coordinates are converted into Allen CCF coordinates by multiplying 1000 and adding 5400 (for AP) and 5700 (for ML). Converted cell coordinates were plotted using Points function in Brainrender v2.0.0.0 by following the instruction (https://github.com/brainglobe/brainrender). Density plots are generated by using kdeplot function in seaborn (https://seaborn.pydata.org).

## Acknowledgment

We thank Lauren McElvain and Harvey J. Karten for discussion on anatomical annotation. We thank Wang lab for helpful discussions and suggestions over the course of this work. This work is support by NIH grants U19 NS107466, and NS 077986.

## Inventory of Supplemental Information

### Supplemental Figures

Figure supplement 1 - related to Figure 1

Figure supplement 2 - related to Figure 2

Figure supplement 3 - related to Figure 5

Figure supplement 4 - related to Figure 5

Figure supplement 5 - related to Figure 5

Figure supplement 6 - related to Figure 5

Figure supplement 7 - related to Figure 5

Figure supplement 8 - related to Figure 7

### Supplemental Movies (available separately online)

Movies can be opened in a Web browser

Supplemental Movie 1 - Related to Figure 5

Interactive Movie: 3D reconstructed whisker, genioglossus, and masseter premotor neurons.

### Source files

Coordinates of premotor neurons from 12 animals.

Coordinates.zip (available upon acceptance)

**Figure supplement 1.**
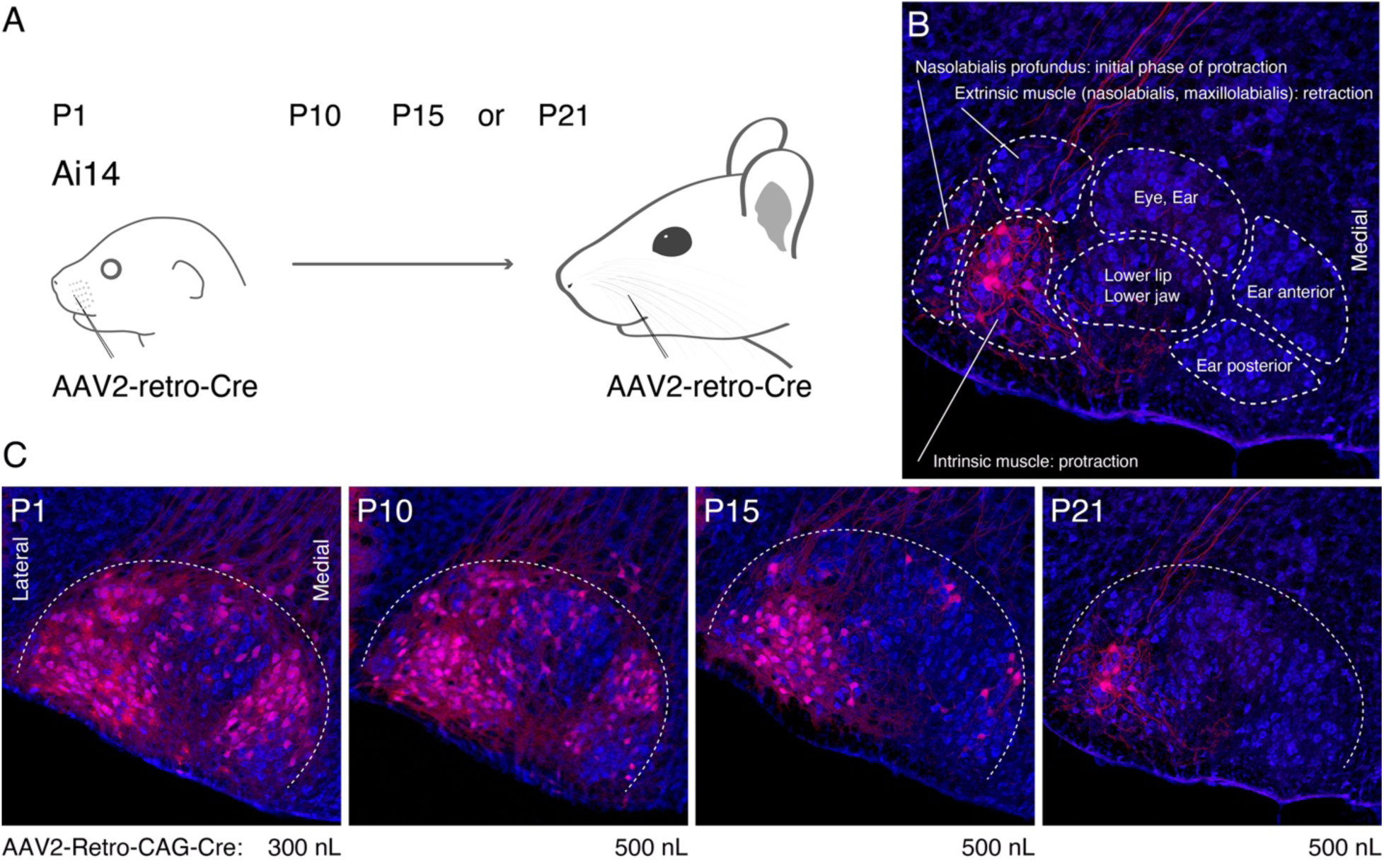
(**A**) Schematic of AAV2-retro-Cre injection. AAV2-retro-Cre is injected into the whisker pad of Ai14 mice at P1, P10, P15, or P21. (**B**) Subnuclei of VII. Each subnucleus is circled by dotted lines. Peripheral muscle targets are shown in each subnucleus. Motoneurons in the lateral part of VII (red) are labeled by AAV2-retro-Cre injection at P21. (C) Labeling patterns from P1, P10, P15, or P21 injected animals. Injection volumes of AAV2-retro-Cre are shown under each panel. Sections were counterstained with fluorescent Nissl (blue).

**Figure supplement 2.**
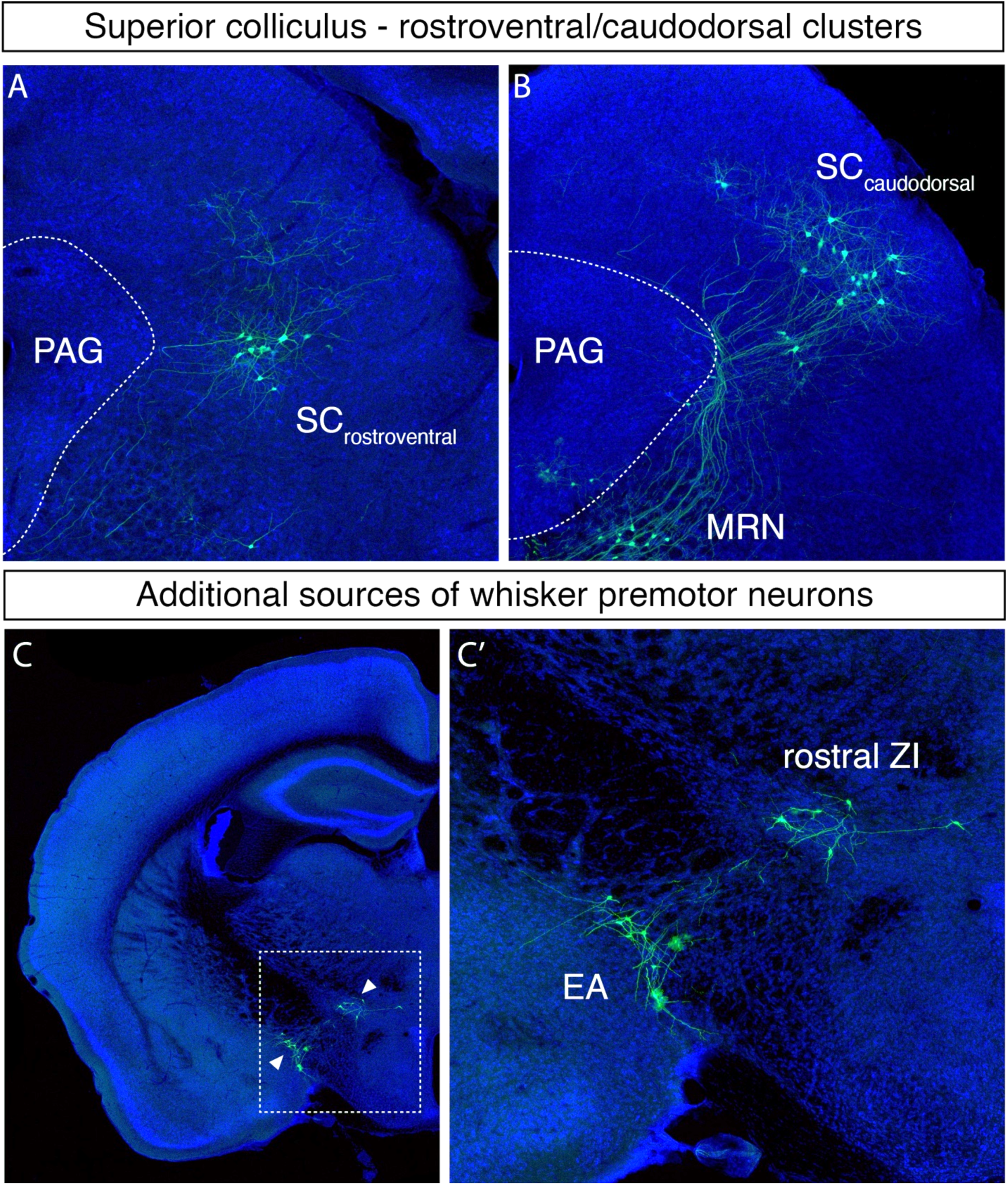
Additional whisker premotor inputs. (**A, B**) Labeled neurons in the rostroventral (**A**) and caudodorsal superior colliculus (**B**). (**C**) Labeled neurons are observed in the rostral zona incerta (ZI) and extended amygdala (EA) (a magnified image of the boxed area is shown in **D’**).

**Figure supplement 3.**
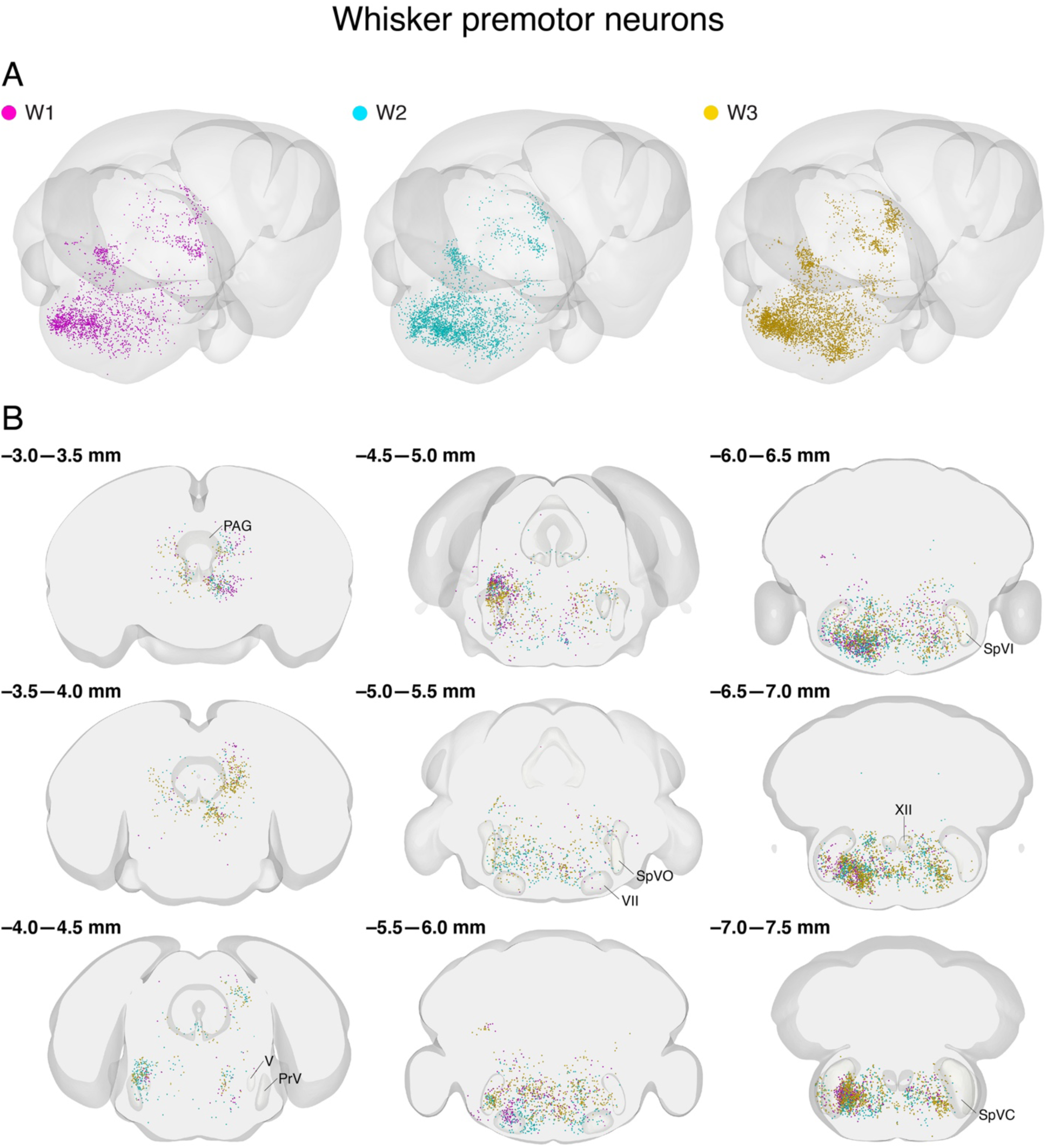
Distribution of labeled whisker premotor neurons from individual animals in Allen CCF. (**A**) 3D reconstruction of labeled whisker premotor neurons with posterior oblique view from 3 different mice (W1 magenta, W2 cyan, W3 gold). (**B**) Coronal views of reconstructed whisker premotor neurons from all three mice in the same coordinates. Anterior-posterior levels (referenced to Bregma) are shown on the top left of each panel.

**Figure supplement 4.**
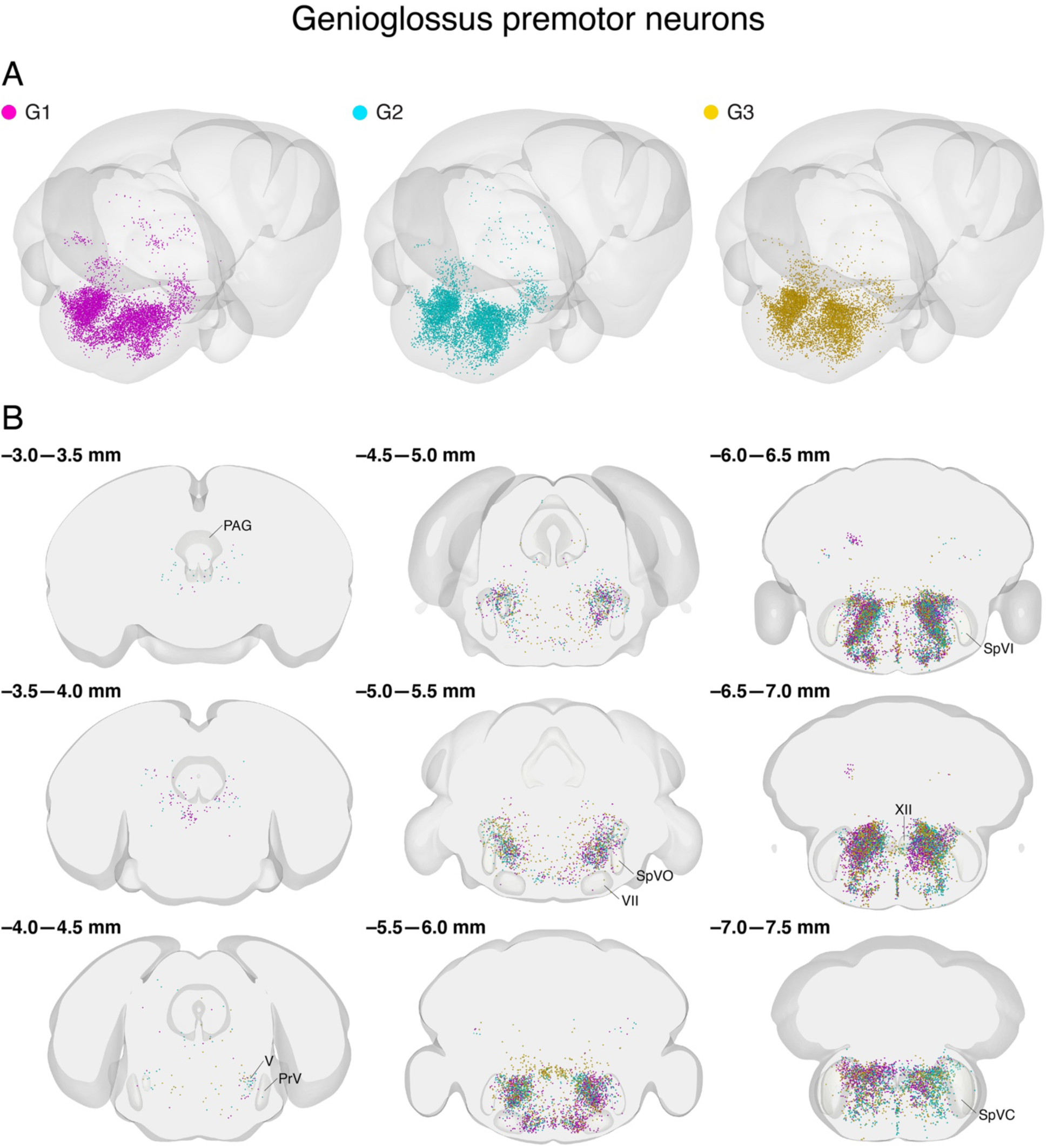
Distribution of labeled genioglossus premotor neurons from individual animals in Allen CCF. (**A**) 3D reconstruction of labeled genioglossus premotor neurons with the posterior oblique view from 3 different mice (G1 magenta, G2 cyan, G3 gold). (**B**) Coronal views of reconstructed genioglossus premotor neurons from all three mice in the same coordinates. Anterior-posterior levels (referenced to Bregma) are shown on the top left of each panel.

**Figure supplement 5.**
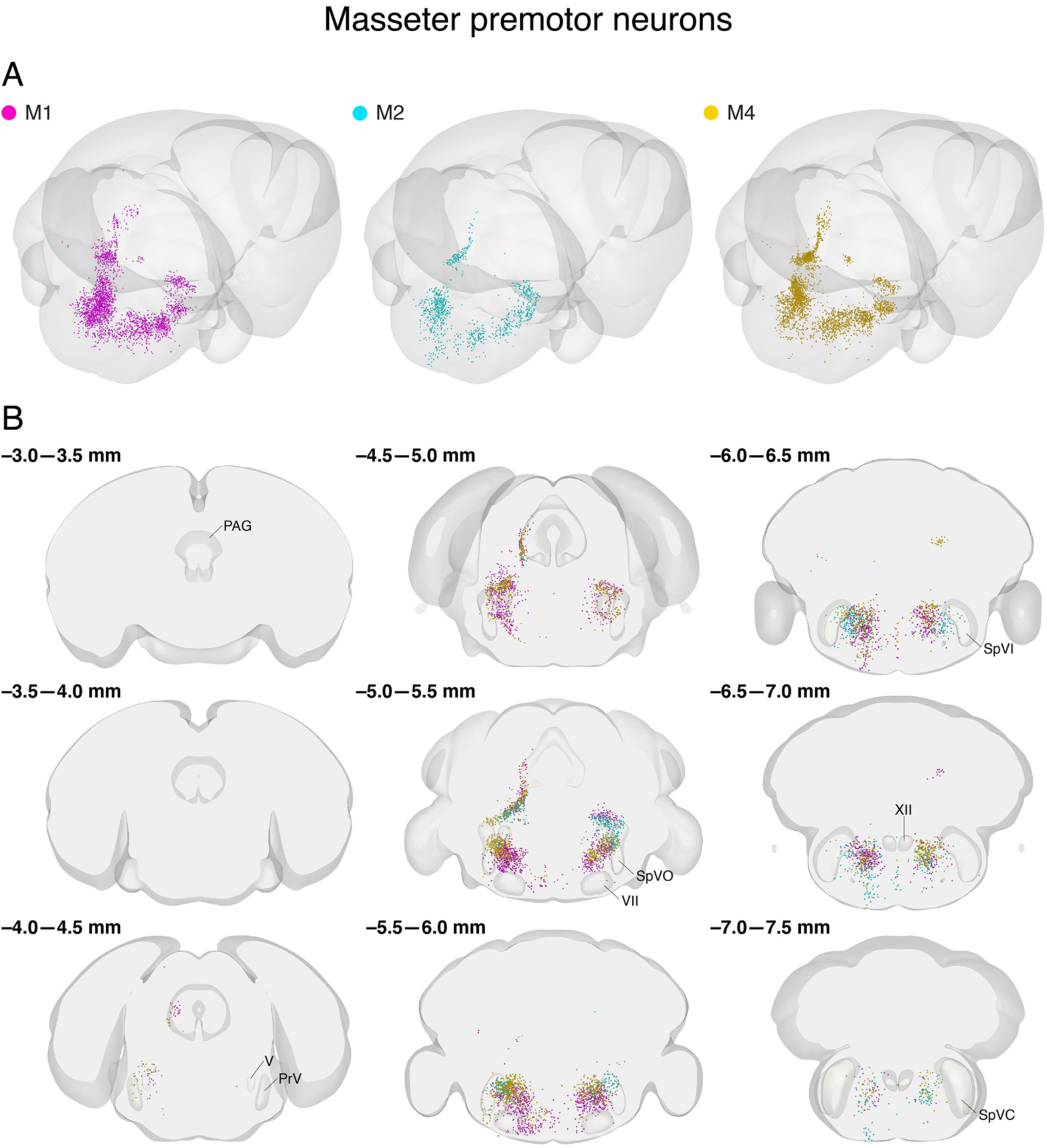
Distribution of labeled masseter premotor neurons from individual animals in Allen CCF. (A) 3D reconstruction of labeled masseter premotor neuron with the posterior oblique view from 3 different mice (M1 magenta, M2 cyan, M3 gold). (B) Coronal views of reconstructed masseter premotor neurons from all three mice in the same coordinates. Anterior-posterior levels (referenced to Bregma) are shown on the top left of each panel.

**Figure supplement 6.**
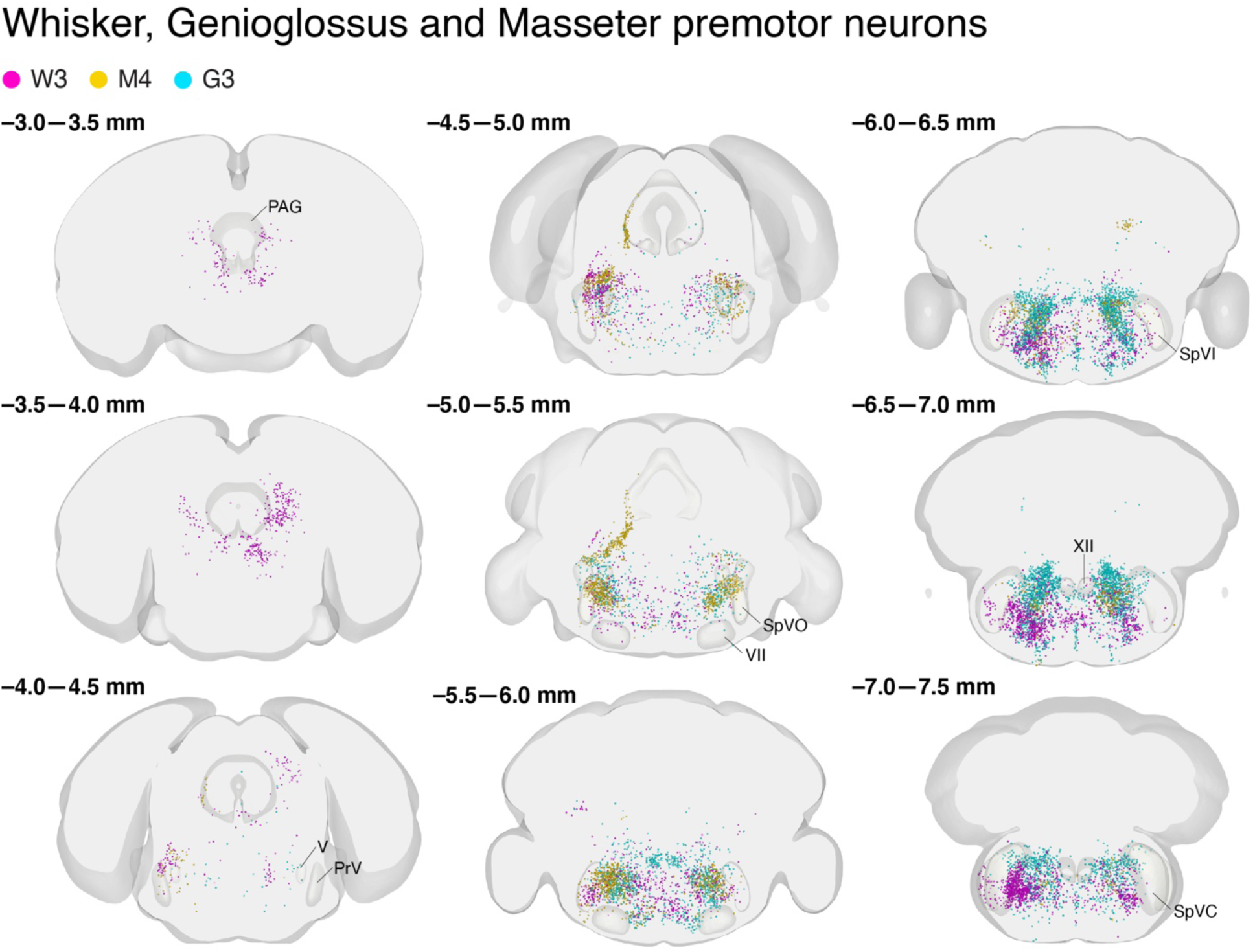
Whisker-, genioglossus-, and masseter- premotor neurons in the same Allen CCF. Coronal views of reconstructed whisker (magenta), genioglossus (cyan), and masseter (gold) premotor neurons. Anterior-posterior levels (referenced to Bregma) are shown on the top left of each panel. The identification numbers of animals are shown on top.

**Figure supplement 7.**
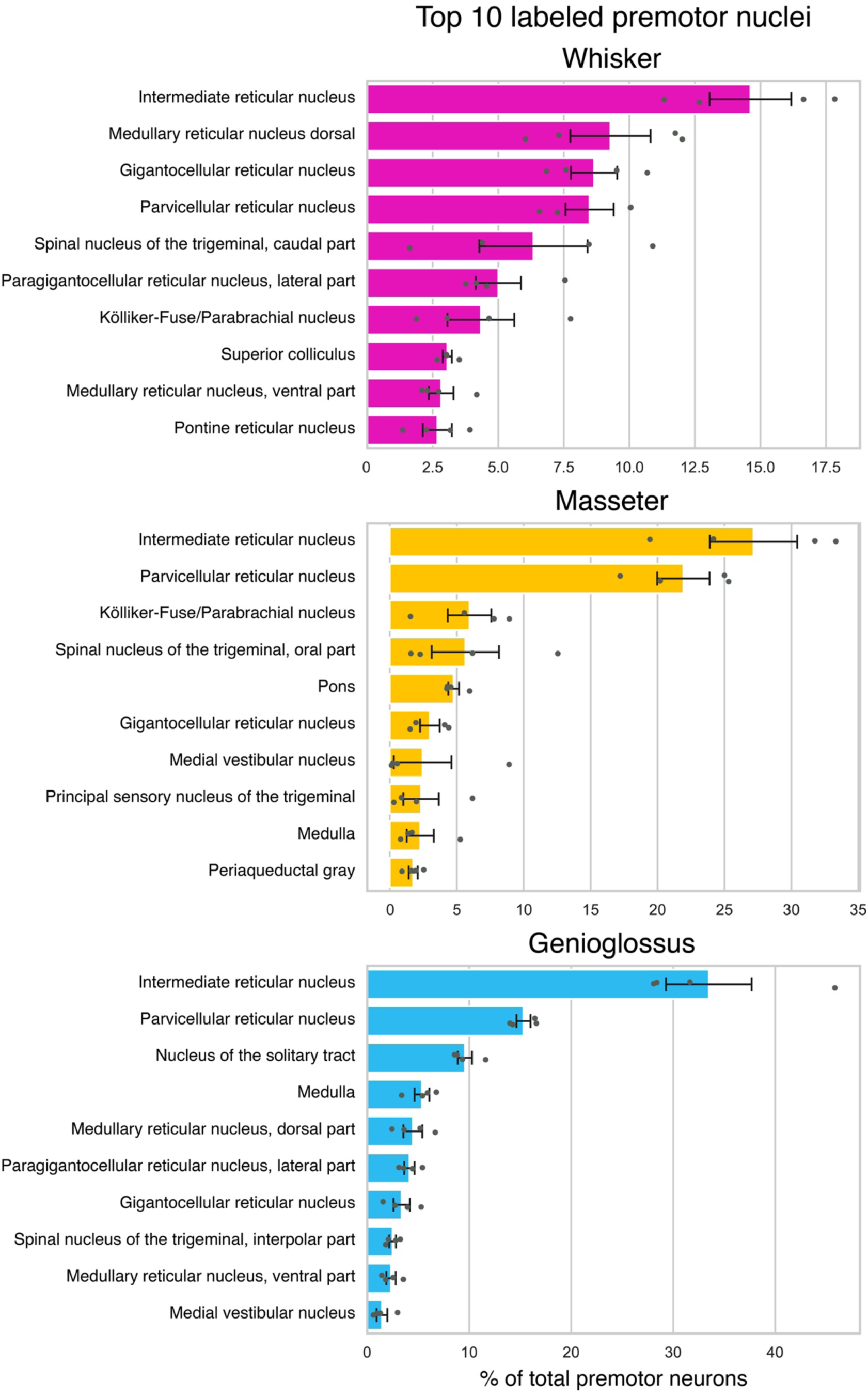
Quantification of trans-synaptically labeled neurons in top 10 labeled brain areas for each motor group based on Allen CCF nomenclature. Summary of the distributions of whisker (Top, magenta, n = 4), masseter (middle, gold, n = 4), and genioglossus (bottom, cyan, n = 4) premotor neurons. Brain areas were automatically annotated based on Allen CCF coordinates. The value is normalized against the total numbers of labeled neurons and averaged across animals. Data are mean ± SEM (n = 4).

**Figure supplement 8.**
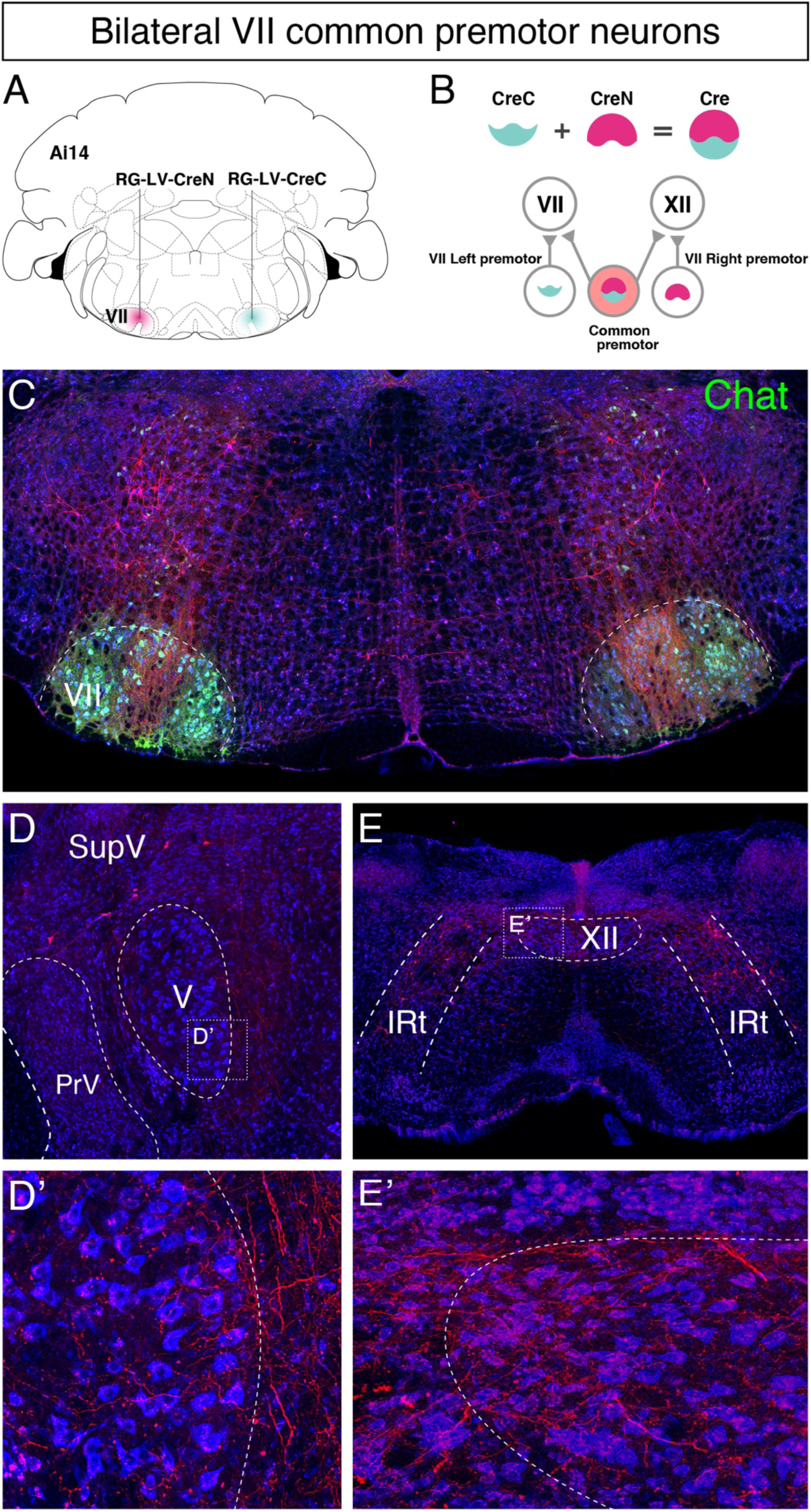
Identifying common premotor neurons with bilateral collateral projections to VII_middle_. (**A, B**) Schematic of split-Cre tracing strategy. (**B**) RG-LV-CreN and RG-LV-CreC were injected into the left and right VII_middle_ of Ai 14 mice, respectively. Cre is reconstituted only in neurons innervating both left and right VII_middle_, which induces tdTomato reporter expression. (**C**) Representative images of axons/axon collaterals in the injection sites. Note the dense tdTomato signal in VII_middle_, and bilateral VII_middle_ common premotor neurons in th IRt dorsal to VII. Motoneurons were stained with anti-chat antibody (green). (**D-E’**) bilateral VII_middle_ common premotor neurons are observed in SupV (D) and the dorsal IRt (E). Axon collaterals of bilateral VII_middle_ premotor neurons also innervate V (**D’**; the boxed area in D) and XII (**E’**; the boxed area in E) motor neurons. Sections were counterstained with fluorescent Nissl (blue).

Supplemental Movie 1. Interactive Movie: 3D reconstructed whisker, genioglossus, and masseter premotor neurons.

Whisker (w3, magenta), genioglossus (g3, cyan), and masseter (m4, gold) premotor neurons were reconstructed in the same Allen CCF. The file can be opened in a Web browser. The 3D reconstructed brain can be rotated by clicking and dragging it. Zoom can be controlled with the mouse wheel.

